# Three Prime Repair Exonuclease 1 preferentially degrades the integration-incompetent HIV-1 DNA through favorable kinetics, thermodynamic, structural and conformational properties

**DOI:** 10.1101/2024.03.19.585766

**Authors:** Prem Prakash, Purva Khodke, Muthukumar Balasubramaniam, Benem-Orom Davids, Thomas Hollis, Jamaine Davis, Jui Pandhare, Bajarang Kumbhar, Chandravanu Dash

**Affiliations:** Department of Biochemistry, Cancer Biology, Neuroscience and Pharmacology, Meharry Medical College, Nashville, Tennessee, 37208, USA; Sunandan Divatia School of Science, NMIMS University, Mumbai, 400056, India; Department of Biochemistry and Molecular Biophysics, Columbia University, New York City, New York, 10032, USA; Department of Biochemistry and Center for Structural Biology, Wake Forest University School of Medicine, Winston-Salem, NC, 27157, USA; Center for AIDS Health Disparities Research, Meharry Medical College, Nashville, Tennessee, 37208, USA; Department of Microbiology, Immunology, and Physiology, Meharry Medical College, Nashville, Tennessee, 37208, USA

## Abstract

HIV-1 integration into the human genome is dependent on 3’-processing of the reverse transcribed viral DNA. Recently, we reported that the cellular Three Prime Repair Exonuclease 1 (TREX1) enhances HIV-1 integration by degrading the unprocessed viral DNA, while the integration-competent 3’-processed DNA remained resistant. Here, we describe the mechanism by which the 3’-processed HIV-1 DNA resists TREX1-mediated degradation. Our kinetic studies revealed that the rate of cleavage (*k*_cat_) of the 3’-processed DNA was significantly lower than the unprocessed HIV-1 DNA by TREX1. The efficiency of degradation (*k*_cat_/K_M_) of the 3’-processed DNA was also significantly lower than the unprocessed DNA. Furthermore, the binding affinity (K_d_) of TREX1 was markedly lower to the 3’-processed DNA compared to the unprocessed DNA. Molecular docking and dynamics studies revealed distinct conformational binding modes of TREX1 with the 3’-processed and unprocessed HIV-1 DNA. Particularly, the unprocessed DNA was favorably positioned in the active site with polar interactions with the catalytic residues of TREX1. Additionally, a stable complex was formed between TREX1 and the unprocessed DNA compared the 3’-processed DNA. These results pinpoint the biochemical mechanism by which TREX1 preferentially degrades the integration-incompetent HIV-1 DNA and reveal the unique structural and conformational properties of the integration-competent 3’-processed HIV-1 DNA.

## Introduction

The Three Prime Repair Exonuclease 1 (TREX1) enzyme is the most active 3′→5′ exonuclease in mammalian cells (1). TREX1 degrades cellular DNA originating from aberrant DNA replication, recombination, and repair, along with the DNA intermediates of endogenous retroelement replication (2,3). Removal of these DNA is essential to prevent innate immune response to self-DNA (2,3). Thus, the primary function of TREX1 is to degrade the immune-stimulatory DNA (ISD) to prevent autoimmune response (4). Accordingly, TREX1 mutations are linked to autoimmune disorders including systemic lupus erythematosus, chilblain lupus, Aicardi-Goutières syndrome (AGS) type 1, and retinal vasculopathy with cerebral leukodystrophy (5–8). Interestingly, there is evidence that viruses such as human immunodeficiency virus-1 (HIV-1) usurp TREX1 as a mechanism to counter the anti-viral effects of the host immune system (9,10).

HIV-1 primarily infects T lymphocytes expressing CD4 receptors and CXCR4/CCR5 co-receptors (11). The viral envelope glycoproteins bind to these receptor/co-receptors to mediate fusion of the viral membrane with the cellular plasma membrane resulting in the release of the viral capsid into the cytoplasm (12). The capsid encases two copies of the single-stranded (ss) viral RNA genome and other viral/host factors that are required for replication (13–17). Particularly, the reverse transcription complex (RTC) converts the ssRNA genome into a double-stranded (ds) DNA (18). Then, the pre-integration complex (PIC) carries out integration of the viral DNA into the host chromosomes to establish a provirus (19–22). The provirus is then transcribed by the cellular transcription machinery to produce the spliced viral mRNAs that encode viral proteins and the full-length unspliced genomic RNA to be packaged into the progeny virions (23). Importantly, these early steps of HIV-1 infection generate a number of viral nucleic acid intermediates such as ssRNA, dsRNA, ssDNA, dsDNA, and DNA/RNA hybrids (18) that can be degraded by TREX1 (1). However, the mechanisms by which TREX1 recognizes and degrades these HIV-1 DNA and RNA substrates in an infected cell to regulate innate immune response is poorly understood.

The role of TREX1 during HIV-1 infection was first reported in a study of the human SET complex (9), that contains TREX1 and two other nucleases (24,25). Knockdown of TREX1 inhibited HIV-1 infection by increasing autointegration of the viral DNA concurrent with reduced proviral integration (26). TREX1 knockdown also resulted in the accumulation of viral DNA and induction of interferon stimulatory genes (ISGs) (4). Conversely, TREX1 overexpression suppressed ISGs, indicating that TREX1 regulated innate immune response to HIV-1 infection (9). Thus, it was predicted that by degrading accumulating viral DNA, TREX1 enables the virus to evade the innate immune response. Recently, we reported that TREX1 expression is elevated in HIV-1 infected cells (27). Higher TREX1 levels correlated with increased proviral integration (27) providing further support for a functional role of TREX1 in HIV-1 infection.

HIV-1 integration is dependent on 3’-processing of the reverse transcribed viral DNA. Specifically, HIV-1 integrase (IN) enzyme cleaves the 3’-GT dinucleotides from both the U5 and U3 ends of the long terminal repeats (LTR) to create the recessed 3’-CA_OH_ ends (Fig. 1A) (20). Then during the strand-transfer step, the recessed 3’-OH group of the processed viral DNA ends carries out nucleophilic attack on the phosphodiester backbone of the chromosomal DNA to insert the viral DNA (20). The requirement of the recessed 3’-OH group for the strand transfer step renders the unprocessed HIV-1 DNA incompatible for proviral integration. Interestingly, we observed that TREX1 preferentially and efficiently degrades the unprocessed HIV-1 DNA ends that are incompetent for integration (27). Surprisingly, the integration-competent 3’-processed viral DNA substrates remained highly resistant to degradation by TREX1 (27). However, the mechanism underlying the preferential degradation of the integration incompetent HIV-1 DNA substrates and the resistance of 3’-processed viral DNA by TREX1 is not fully understood.

**Figure 1.**
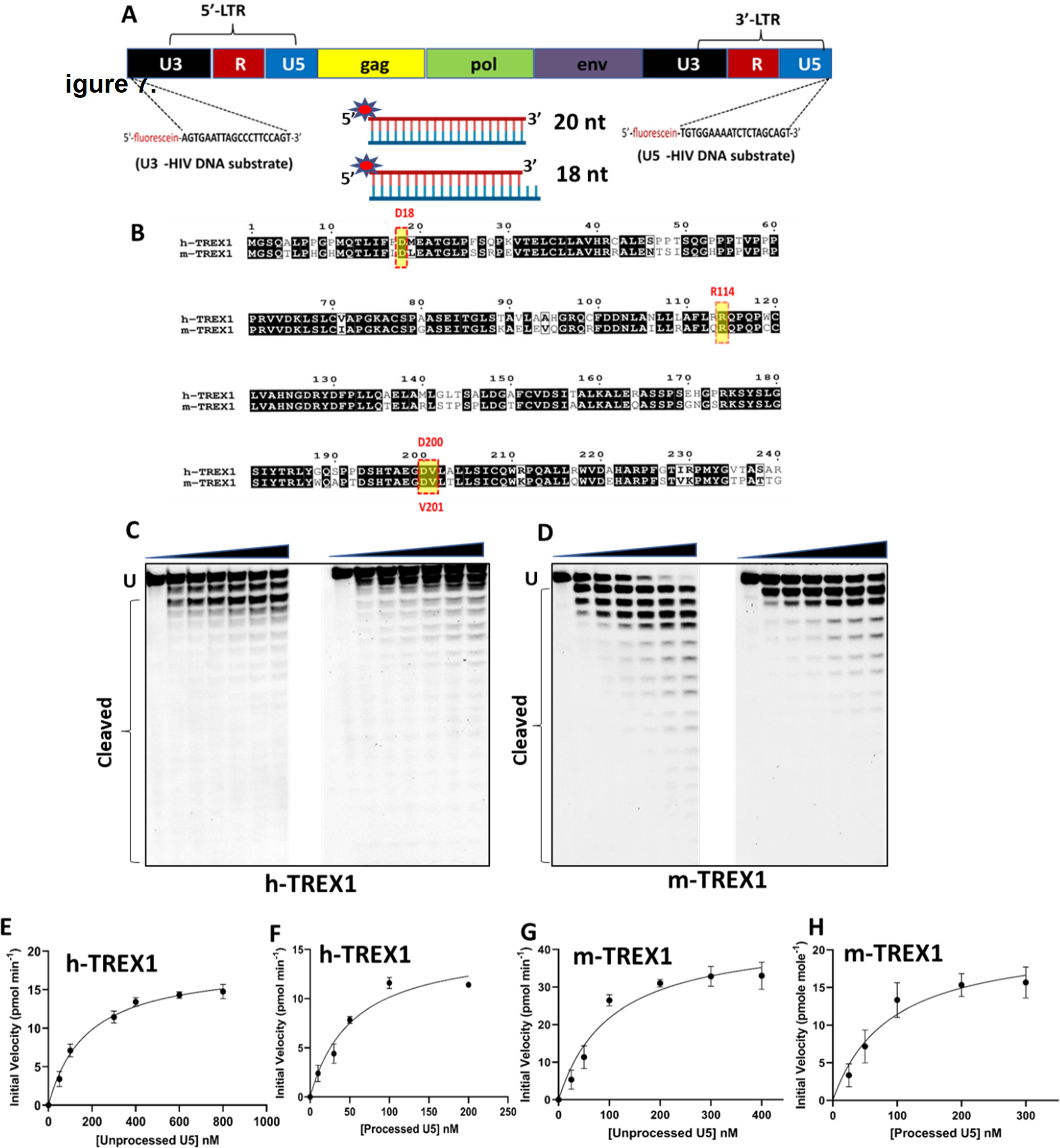
TREX1-mediated degradation of U5 viral DNA substrates. **(A)** Design of DNA substrates containing the U5 and U3 sequences of the HIV-1 LTR. The schematic representation of HIV-1 genome with the LTR consisting of U5 and U3 ends shown in black and blue colors respectively. The 20 nucleotides sequence from U3 and U5 ends used as the viral DNA substrates to determine the kinetic parameters are indicated by dashed lines at both the ends. Schematics of the dsDNA substrates mimicking the blunt-ended unprocessed strand and the 3’-processed strand is also shown. **(B)** Amino acid sequence comparison of human TREX1 (h-TREX1) and mouse TREX1 (m-TREX1). The highlighted active site residues are highly conserved in both the species. **(C-D)** Representative sequencing gel images showing the activity of h-TREX1 **(C)** and m-TREX1 **(D)** with unprocessed (left panel) and 3’-processed (right panel) U5 HIV-1 DNA substrates. Uncleaved (U) and cleaved substrates are marked in these gels. The uncleaved and cleaved products of both unprocessed and processed viral DNA substrates respectively in panels C and D were separately analyzed and aligned at the same level for the purpose of comparison. The kinetic progression curve for the **(E-F)** h-TREX1 and **(G-H)** m-TREX1 activity with U5 unprocessed and U5 3’-processed HIV-1 DNA substrates. Each result and the data points on the kinetic progression curves are the representative of three experimental replicates with errors bars representing SEM.

In this study, we describe the biochemical mechanism that governs the selectivity of TREX1 activity to the blunt-ended unprocessed HIV-1 DNA compared to the 3’-processed DNA. We performed TREX1 activity measurements to calculate the kinetic parameters using viral DNA substrates containing the U5 and U3 end sequences of HIV-1 LTR. We found that the rate of cleavage (*k*_cat_) of the unprocessed viral DNA by TREX1 was higher than that of the 3’-processed viral DNA. Moreover, the catalytic efficiency (*k*_cat_/K_M_) of TREX1 for the unprocessed viral DNA was significantly higher than the 3’-processed viral DNA. In fluorescence polarization anisotropy analysis, we observed that TREX1 exhibited marked differences in the binding affinity (K_d_) for unprocessed and processed viral DNA substrates. To further identify the biochemical and structural determinants, we performed molecular docking and molecular dynamics simulation studies of TREX1 with the viral DNA substrates. Results from these *in silico* studies revealed novel conformational differences in the binding of unprocessed and processed viral DNA substrates to the TREX1 active site. These studies also identified the formation of a stable complex between TREX1 and the unprocessed viral DNA compared to the complex with the processed viral DNA. Collectively, our results provide kinetic, thermodynamic, and structural insights into the preferential degradation of integration-incompetent HIV-1 DNA by TREX1.

## Materials and Methods

### Purification of TREX1

To measure the exonuclease activity of TREX1 (mouse and human TREX1), a truncated N-terminal (1-242 amino acids) of TREX1 containing the catalytic domain that displays exonuclease activity was overexpressed and purified. The plasmid construct used to express the human TREX1 enzyme in *Escherichia coli* and purification of the TREX1 used for structural studies has been described previously (28–34), with few variations. The TREX1 gene is expressed as fusion with maltose binding protein (MBP) in pLM303. The constructs for mouse-TREX1 (m-TREX1) and human-TREX1 (h-TREX1) were transformed into Rossetta II strain of *Escherichia coli* cells and plated on selective media containing Kanamycin (Kan) and Chloramphenicol (CP). Using a colony from the transformation plate, inoculate a ∼ 200 mL LB/Kan/CP broth and then determined the absorbance A_600_ of the starter culture. Then, the starter culture was diluted into five 1 L broths such that their A_600_ ∼ 0.1. The new cultures were then incubated at 37°C/180 rpm and the absorptions at A_600_ were periodically measured, until A_600_ = 0.5–0.8. After 2 hours, each culture was induced with 0.5 M IPTG (Sigma Aldrich). The culture was placed back in incubation at 37°C/180 rpm for 10 min. Then, the cultures were retrieved and cooled in ice baths until their temperature reaches 20°C as measured using a thermometer. After cooling, cultures were placed into incubator at 16°C/180 rpm to incubate overnight (15–20 hours). Induced cultures were then centrifuged at 4000 rpm/4°C for 20 min. The supernatants were then discarded, and the obtained pellet was resuspended into 120 mL Amylose column A Buffer (20 mM Tris-base-HCl (pH 7.5), 200 mM NaCl, 200 mM NaCl) +10% glycerol, and three protease inhibitor tablets (Calbiochem set III). The cells were lysed using ultrasonication (Branson analogue sonifier 450). The lysate was centrifuged for 20 min at 15,000 rpm/4°C and the supernatant was collected. All purification steps were performed at 4°C and the pre-packed amylose column (5ml Hi-Trap GE healthcare) was equilibrated with 25 mL Amylose column A Buffer. The cleared lysate sample was loaded to the column using a flow rate of 1.5 mL/min. The column was then washed with 120 mL Amylose A Buffer and the elution of protein from column with Amylose B buffer (Amylose column buffer A + 10 mM Maltose) was performed with an addition of final 10% glycerol to the eluate. The entire purification with amylose column was carried out using fast performance liquid chromatography (Bio-Rad NGC). Dialysis of the protein was performed for 2 hours in 1 L of P-cellulose column A buffer (50 mM Tris-base-HCl (pH 7.5), 1 mM EDTA, 10% (v/v) glycerol containing 50 mM NaCl), using 6–8 kDa MWCO (Millipore, Novagen) dialysis tubes was carried out at 4°C. The dialyzed sample was loaded into the manually packed 15 ml Phospho-cellulose (Creative Biomart) column. The column was then washed with 75 mL P-cell A Buffer. The elution was then carried out using step gradient of 0-100% P-cell B buffer with 2 ml fraction volumes and combined the eluent fractions containing TREX1. The concentrations of purified TREX1 were determined by absorption of A280 using the molar extinction coefficient ɛ = 23,950 M^−1^ cm^−1^, and the purity was confirmed using SDS PAGE.

### Exonuclease activity and kinetics measurement of TREX1

TREX1 activity was measured using previously described method (30). Exonuclease assays were performed with either 18-mer or 20-mer fluorescein-labeled oligonucleotides annealed with unlabeled complementary 20-mer oligonucleotides (Fig. 1A). The sequences of the oligonucleotides are: 20-mer U5 (unprocessed), 5’-fluorescein-TGTGGAAAATCTCTAGCAGT-3’; 18-mer U5 (processed), 5’-fluorescein-TGTGGAAAATCTCTAGCA-3’; 20-mer U5 (complement), 5’-ACTGCTA GAGATTTTCCACA-3’. 20-mer U3 (unprocessed), 5’-fluorescein-AGTGAATTAGCCCTTCCAGT-39; 18-mer U3 (processed), 5’-fluorescein-AGTGAATTAGCCCTTCCA-3’; 20-mer U3 (complement), 5’-ACTGGAAGGGCTAATTCACT-3’. The dsDNA substrates were generated by annealing the fluorescein labeled 20-mer with unlabeled complementary 20-mer oligonucleotide. Annealing was done by mixing 10 µM 5’-fluorescein-labeled oligonucleotide with 20 µM unlabeled complementary oligonucleotide in annealing buffer (10 mM Tris-Cl pH 7.5 and 10 mM NaCl) and then heating at 95°C for 5 min. The annealing mixture was slowly cooled to the room temperature. The annealed double stranded DNA substrates were confirmed with 20% native PAGE.

Exonuclease assays were carried out in a 25 µl reaction mixture. The reaction buffer consisted of 20 nM Tris-Cl (pH 7.5), 5 mM MgCl_2_, 2 mM DTT, 100 µg/ml bovine serum albumin double stranded DNA substrates (10 nM to 1 µM) and required concentration of TREX1 enzyme. The substrate and reaction buffer were preincubated at 37°C and then the enzyme was added to start the reaction. The reactions were stopped using loading buffer (20 mM EDTA, 76% deionized formamide and 10% xylene cyanol) at different time points followed by heating at 95°C for 2 min. Each reaction was carried out at various time points for the kinetics measurement with an extra no-enzyme control reaction. Exonuclease reaction products were then resolved on a 23% 29:1 acrylamide/bis-acrylamide 7 M urea PAGE gel. The cleaved products were visualized using PharosFX plus molecular imager (Bio-Rad laboratories) and the band intensities were quantified and analyzed using Quantity One 1-D software (Bio-Rad Laboratories).

TREX1 activity was quantified by measuring the band intensities in each lane of the (reaction) and summed called lane intensity (30,32,35). The lane intensity corresponds to the total fluorescein labeled used in the reaction (e.g., 100 nM in 25 µl reaction mixture, contains 2.5 pmol of fluorescein-labeled oligonucleotide). The ratio of band intensity to lane intensity when multiplied by the number of moles of fluorescein-labeled oligonucleotide yields the amount (moles) n-mer oligonucleotide as each band in the lane. The top band is of the highest molecular weight and each band below represents the degraded product of length decreased by a single nucleotide. The total number of moles of dNMPs excised in each reaction (lane) is calculated by multiplying the moles of each cleaved oligonucleotide by the number of excised n^th^-mer oligonucleotide associated with each band and then these values for all the bands were summed up for a given lane. To calculate the excised dNMPs in each reaction (lane) are as follows:

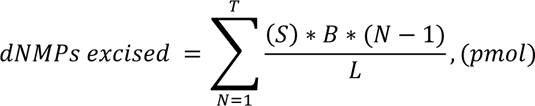

Where, S= Total Number of moles of DNA substrate in the reaction mixture, N= Band number in the given lane, B= Band intensity associated with N^th^ band, L= Lane intensity of each reaction (sum of B and N in the given lane), and T= Total number of bands in the given lane. Once the amount of excised dNMPs calculated at different time points, the number of products (pmol) were plotted as a function of time and the slope of the linear curve represented the initial velocity (pmol min^−1^ ml^−1^) of TREX1 with corresponding substrate concentration. Then initial velocities of TREX1 (at fixed concentration) were calculated for varying the substrate concentrations that yields the saturation hyperbolic curve. The curves were fitted to the non-linear regression model of Michaelis Menten equation using Graph Pad software. The kinetics parameters of TREX1 exonuclease activity such as maximum velocity (V_max_), Michaelis constant (K_M_) and catalytic turnover number (*k*_cat_) for each substrate were calculated.

For steady-state kinetic investigation, the relation between K_M_ and K_d_ were calculated as per the following formulas (36).

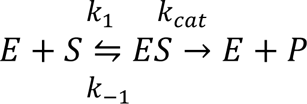

Here, *E* = [enzyme], *S* = [Substrate], P = [Product], *k*_1_: the association rate constant of *ES* – complex (M^−1^ s^−1^), *k_-_*_1_: the unproductive dissociation rate constant of *ES* – complex (s^−1^), *k*_cat_: the turnover number (s^−1^). So, according to steady state approximation,

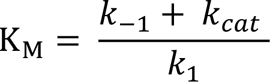

Here, K_M_ is the Michaelis constant. The Michaelis-Menten approximation for the rapid equilibrium for an enzymatic reaction as shown above is valid only if k_-1_ >> *k*_cat_.

Now since,

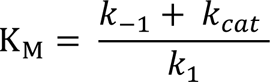

Then,

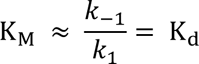

To establish a relationship between K_M_ and K_d_, the rapid equilibrium states that the dissociation rate constant k_-1_ should be fast in comparison to the turnover number (*k*_cat_). Thus, we can assume that the substrate concentration at the half maximal velocity could be equal to the dissociation constant K_d_.

### Fluorescence Polarization Anisotropy (FPA) assay

Fluorescence anisotropy assay were carried out to measure the binding affinity constant (K_d_) of TREX1-DNA complexes, using commercially synthesized fluorescein-labeled oligonucleotides (Thermo Fisher, Waltham, MA, USA). Because TREX1 starts catalyzing the degradation of DNA substrate once the conducive buffer condition is provided, we used CaCl_2_ instead of MgCl_2_ as Mg^2+^ is the cofactor of TREX1. By doing so, the catalysis could be halted but the possibility of binding may not be affected (28). Measurement of direct binding of DNA substrates to TREX1 were carried out by the increase in fluorescence polarization (FP) upon titration of a fixed concentration of the DNA substrates with purified TREX1. The titration was performed in a black 96 well plate (Thermo-Fisher scientific) that allows the measurement of multiple data points and facilitating the titration curve. FP versus TREX1 concentration were plotted to generate a saturation curve that begins at the free DNA substrate base line polarization. The substrate was then titrated by varying the concentration of TREX1 (4 µM) which was then diluted two-fold from top to the consecutive wells, each having half the concentration of preceding ones. The concentration of unprocessed and processed DNA substrates was kept constant at 10 nM. After addition of the DNA substrates to the enzyme, the reaction mixtures were incubated at 4°C for 1 hour. The fluorescence polarization measurement for the binding of TREX1 and DNA substrates were carried out using an optical system that includes polarizing filters in the light path. The filters used in these experiments were of excitation/emission wavelength of 485/530 nM. Samples in the microplates are excited, and depending on the mobility of the fluorescent molecules, the polarization of emitted light changed and that was monitored and analyzed using in the Gene5 data analysis software. The increasing polarization values were plotted against enzyme concentration and the progression curve were fit into the one site specific binding equation of Graph Pad Prism software (version 9.0).

### Molecular docking studies

Molecular docking of TREX-1 (PDB ID:7TQQ) with double stranded viral DNA end substrates (U5 and U3 ends) were performed using the HDOCK server (http://hdock.phys.hust.edu.cn/), which uses an *ab initio* free docking that is based on hybrid algorithm of template-based modeling (37). The models and coordinates for double stranded DNA oligonucleotide substrates were generated using COOT (38). HDOCK samples all possible binding modes between protein and the ligand by using a fast Fourier transform based algorithm (39). The sampled modes of binding are then evaluated through iterative knowledge-based scoring function (40). The finalized binding mode of protein-DNA interactions was evaluated by rank and docking scores. The protein-DNA complex structure models with lowest root-mean-square deviation (RMSD) and highest docking score were selected for analysis of the binding modes. Further, the interacting key residues of the three-dimensional model of protein-DNA complexes were examined through COOT (38). The non-bonded interactions such as electrostatic, van der Waals, hydrophobic, hydrogen bonds interactions in the protein complex models were determined using protein interaction calculator (41). The putative interfacial interactions of TREX1-DNA complex structures were evaluated using PDBePISA (Protein, Interface, Structures and Assemblies) server (42). Final representation of protein DNA complexes were done by using software PyMol (http://www.pymol.org).

### Molecular dynamics simulation

Molecular dynamics (MD) simulations were performed using Gromacs2021.5 (43) to explore the binding mode and interaction of human TREX1 with U3 and U5 processed and unprocessed DNA substrates. The least binding energy docked conformation of TREX1 with U3 and U5 processed and unprocessed DNA substrates, respectively were used as starting conformation for molecular dynamics simulation. For the MD simulation, amber ff99SB force field were used. All the simulation systems were solvated using a TIP3P water module in a cubic box of 10 Å, and the required number of counter ions was then added to bring the system to a neutral state using ‘xleap’ module of AmberTools18 (44) Using the ‘Parmed tool’, the amber ‘topology’ and ‘co-ordinate’ files were transformed into the companion ‘top’ and ‘gro’ files for Gromacs for simulation (https://github.com/ParmEd) similar to an earlier study (45). For all the simulated complexes, the steepest descent (5000 steps) and the conjugate gradient (2000 steps) methods were used to do the energy minimizations. To equilibrate all the systems, 1 ns NVT and 1 ns NPT simulations were performed. Subsequently, the production MD simulation of 400 ns was performed for each system using the cut-off distance of 1.0 nm with a Fourier spacing of 0.16 nm and an interpolation order of 4 was used, and the particle mesh ewald (PME) method was used to calculate the long-range electrostatic interactions (46). The simulated trajectories and snapshots were further analyzed and visualized using the Visual Molecular Dynamics (VMD) (47) and PyMol software (48), to explore the binding mode and interactions.

### Statistical analysis

All experiments were conducted at least three times with triplicates. Data were expressed as mean ± standard error of mean (SEM) obtained from three independent experiments. Significance of differences between control and treated samples were determined by Student’s *t-*test. The difference between groups were determined by paired, two-tailed student’s t-test. A p-value of < 0.05 was considered statistically significant.

## Results

### Substrate Design

HIV-1 integration is dependent on 3’-processing of the viral DNA ends by the IN enzyme (20). We recently reported that the unprocessed HIV-1 DNA ends are preferentially degraded by TREX1, but the integration-competent 3’-processed DNA ends remained relatively resistant (27). To understand the mechanism by which the 3’-processed HIV-1 DNA resist degradation by TREX1, we planned kinetic studies of TREX1 activity using oligonucleotide substrates containing the sequences of the U5 and U3 ends. The oligonucleotides were chemically synthesized with a fluorescein tag at the 5’-end (Fig. 1A). To prepare the unprocessed DNA substrates, a 20-mer complementary oligonucleotide was synthesized, whereas for the 3’-processed DNA substrates an 18-mer oligonucleotide lacking the GT-dinucleotides from 3’-end was used. The 5’-fluorescein tagged 18-mer or 20-mer oligonucleotides were annealed to the complementary 20-mer oligonucleotide to prepare the DNA substrates. Thus, the 3’-processed substrates contained a recessive DNA end, whereas the unprocessed substrates remained blunt ended. Being a 3’-exonuclease, TREX1 can degrade both 3’-ends of DNA substrates (49,50). We have observed that blocking TREX1 degradation of one strand with a 3’-phosphorothioate modification did not affect the preference of the unprocessed DNA ends (51). However, viral DNA substrates containing a 3’-phosphorothioate modification in one strand are degraded slowly by TREX1 (51). Therefore, we designed unmodified substrates for measuring kinetic parameters in a biochemically and physiologically relevant conditions. The substrates were then subjected to TREX1 activity individually to measure the steady state kinetic parameters such as Michaelis-Menten constant (K_M_), maximum velocity of enzyme activity (V_max_), catalytic constant or the turnover number (*k*_cat_) and the catalytic efficiency (*k*_cat_/K_M_). These kinetic parameters were measured using both the human-(h-TREX1) and mouse-TREX1 (m-TREX1) enzymes. The rationale being that the catalytic residues of both the enzymes are highly conserved (Fig. 1B) and the structural details of m-TREX1 is extensively studied compared to the h-TREX1 (28,52,53).

### TREX1 degrades the unprocessed U5 and U3 HIV-1 DNA at faster rate than the 3’-processed DNA

3’-processing of both U5 and U3 ends of the HIV-1 LTR are critical for the strand transfer step of integration (20). Therefore, we carried out exonuclease activity assays using the unprocessed and 3’-processed substrates containing sequences of both U5 and U3 DNA ends (Fig. 1A). First, we used a range of TREX1 concentration in our assay to identify the optimal enzyme concentration required for the kinetic studies. We determined that 2 nM concentration of h-TREX1 and m-TREX1 is appropriate for measuring the steady state kinetic parameters. Then, we subjected the viral DNA substrates to the activity of h-TREX1 and m-TREX1 separately to calculate the kinetic parameters. A range of viral DNA substrate concentrations (10 nM to 1 µM) were used to measure TREX1 activity as a function of time. The cleavage products were electrophoretic resolved under denaturing conditions, visualized by phosphor-imaging and the band intensities were quantified to calculate kinetic parameters.

We observed that the unprocessed U5 sequences were degraded efficiently by both the h-TREX1 and m-TREX1 when compared to the 3’-processed strands, in accordance with our published studies (51). For instance, TREX1 efficiently degraded majority of the 20-mer unprocessed substrates (Fig. 1C-D, left panels), whereas the 3’-processed DNA ends remained largely resistant (Fig. 1C-D, right panels). Then, time-dependent degradation of unprocessed and 3’-processed U5 substrates were analyzed to estimate initial velocities of the catalytic activity (Suppl. Figs. 1-2). The initial velocity was plotted vs substrate concentrations (Fig. 1E-H) to calculate V_max_, K_M_ and *k*_cat_ using the Michaelis-Menten equation (Table 1). Based on these kinetic analyses, the calculated turnover rate (*k*_cat_) of the unprocessed U5 viral DNA by h-TREX1 was 10.2 s^−1^. By contrast, the *k*_cat_ value of 3’-processed DNA was 3.8 s^−1^, indicating a ∼2.5-fold slower degradation rate of the 3’-processed substrates compared to the unprocessed substrate (Table-1, Fig. 2A). Furthermore, calculation of the catalytic efficiency (*k*_cat_/K_M_) revealed that the unprocessed substrate is degraded by h-TREX1 at a significantly higher efficiency of ∼103.1 pM^−1^s^−1^ compared to ∼70.2 pM^−1^s^−1^ for the 3’-processed substrates (Table-1, Fig. 2B). Similar to the kinetic parameters of the h-TREX1, both the *k*_cat_ and *k*_cat_/K_M_ values of m-TREX1 for the unprocessed substrate were significantly higher than the 3’-processed substrates (Table-1, Fig. 2A-B). The turnover number (*k*_cat_) of m-TREX1 for the unprocessed substrate was 9.5 s^−1^ while that of 3’-processed substrate was 4.1 s^−1^. The *k*_cat_/K_M_ value for the unprocessed substrate was 86.7 pM^−1^s^−1^ compared to 45.2 pM^−1^s^−1^ for the 3’-processed substrate. A comparative analysis of the h-TREX1 and m-TREX1 kinetic parameters indicated that the overall turnover rate of U5 substrates (both unprocessed and 3’-processed) were similar (Fig. 2A) other than the lower catalytic efficiency of m-TREX1 for the U5 3’-processed substrate (Fig. 2B). Collectively, these analyses demonstrate that both h-TREX1 and m-TREX1 degrade the integration-incompetent unprocessed U5 viral DNA ends at a significantly faster rate than the integration-competent 3’-processed U5 viral DNA.

**Figure 2.**
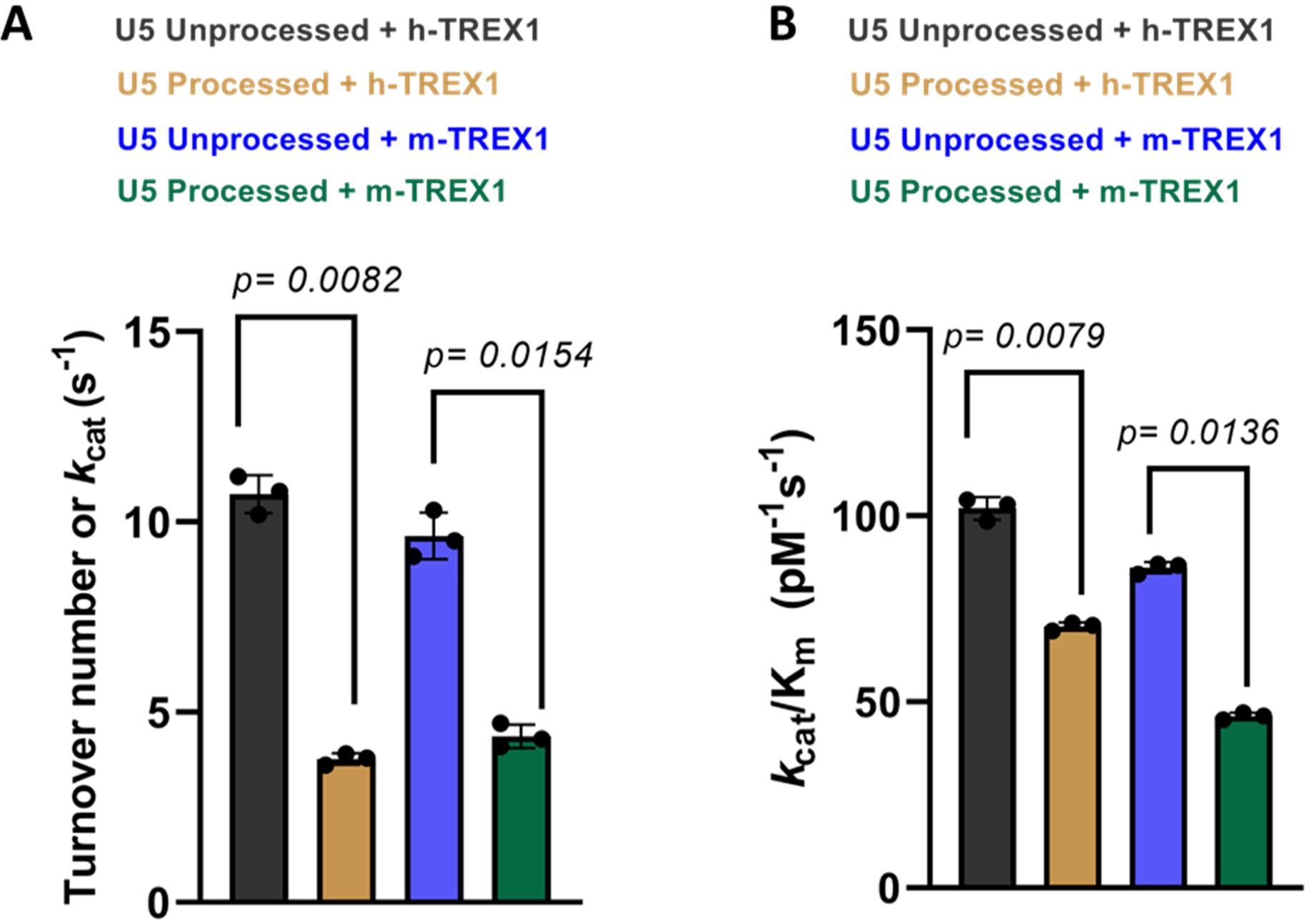
Comparative analysis of the catalytic rate of TREX1 activity with U5 DNA substrates. The representative bar graphs show the quantitative comparison of the rate (*k*_cat_) of h-TREX1 and m-TREX1 activity with U5 **(A)** unprocessed and **(B)** 3’-processed HIV-1 DNA substrates. SEM values + /− from three independent experiments are shown. The p-values are shown just above the bar graphs show statistical significance.

**Table 1.**
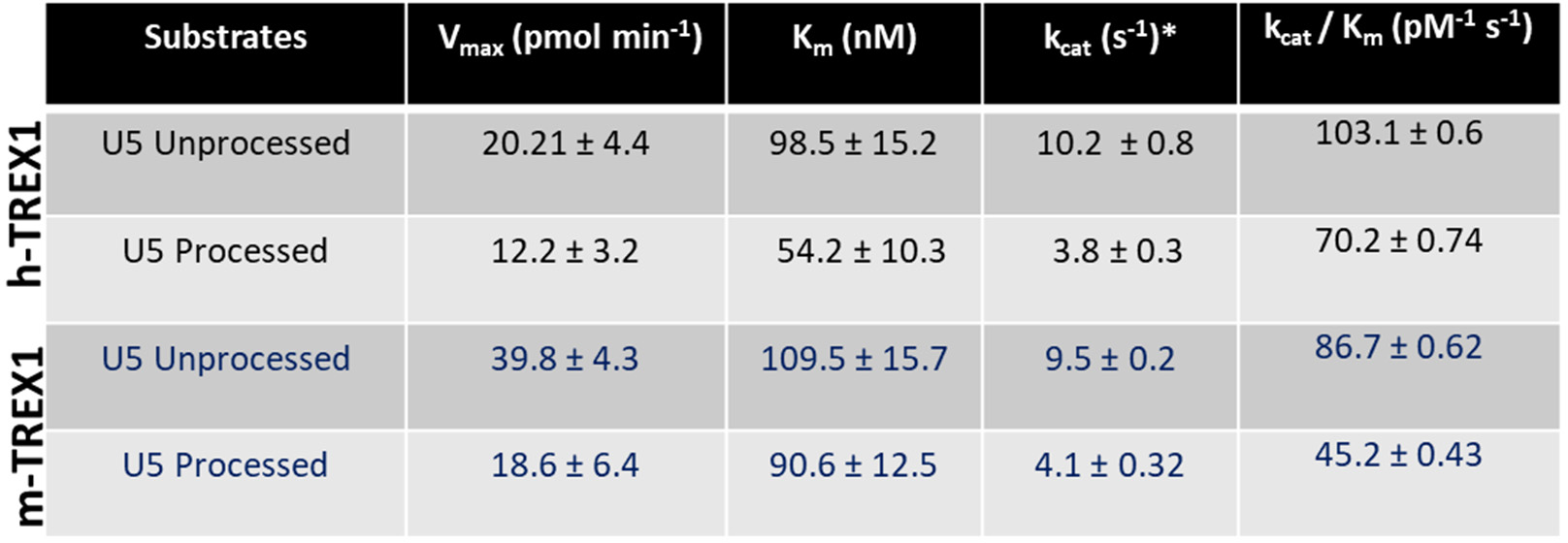

| | Substrates | $V_{\max}$ (pmol min<sup>-1</sup>) | $K_m$ (nM) | $k_{\text{cat}}$ (s<sup>-1</sup>)* | $k_{\text{cat}} / K_m$ (pM<sup>-1</sup> s<sup>-1</sup>) |
| --- | --- | --- | --- | --- | --- |
| h-TREX1 | U5 Unprocessed | 20.21 ± 4.4 | 98.5 ± 15.2 | 10.2 ± 0.8 | 103.1 ± 0.6 |
|  | U5 Processed | 12.2 ± 3.2 | 54.2 ± 10.3 | 3.8 ± 0.3 | 70.2 ± 0.74 |
| m-TREX1 | U5 Unprocessed | 39.8 ± 4.3 | 109.5 ± 15.7 | 9.5 ± 0.2 | 86.7 ± 0.62 |
|  | U5 Processed | 18.6 ± 6.4 | 90.6 ± 12.5 | 4.1 ± 0.32 | 45.2 ± 0.43 |

Next, we measured the kinetics of degradation of the unprocessed and 3-processed substrates containing the U3 ends of HIV-1 DNA (Fig. 3). Similar to the U5 substrates (Fig. 1), degradation of U3 DNA substrates were analyzed at a concentration and time dependent manner (Suppl. Figs. 3-4). The initial velocities were plotted vs substrate concentrations to calculate the kinetic parameters (Fig. 3C-F). Our results showed that the unprocessed U3 substrates were also degraded at a considerably faster rate than the 3’-processed substrates (Fig. 4, Table-2). The *k*_cat_ value of the unprocessed U3 substrates by h-TREX1 was 9.8 s^−1^ (Table 2, Fig. 4A). However, the *k*_cat_ value was 4.5 s^−1^ for the 3’-processed U3 viral DNA, demonstrating a significantly lower (∼2-fold) degradation rate than with the unprocessed DNA. The catalytic efficiency (*k*_cat_/K_M_) of the unprocessed U3 substrate was 63.3 pM^−1^ s^−1^, which was also significantly higher (∼2-fold) than the processed U3 substrates of 28.05 pM^−1^ s^−1^. Accordingly, the kinetic parameters of m-TREX1 showed a higher *k*_cat_ for the unprocessed U3 substrates compared to the 3’-processed substrates (Fig. 4b, Table 2). The turnover number for m-TREX1 with unprocessed substrates was 12.6 s^−1^, while the value was 6.3 s^−1^ for the processed viral DNA. The ratio of *k*_cat_/K_M_ for m-TREX1 degradation of unprocessed substrate (127.8 pM^−1^ s^−1^) was also ∼2.5-fold higher than the 3’-processed U3 viral DNA substrate (54.6 pM^−1^ s^−1^). Interestingly, comparison of the turnover rates and catalytic efficiency between the two enzymes (Fig. 4A-B) suggest that the U3 3’-processed substrate remained resistant to both the enzymes relative to the unprocessed strand. Together, these results clearly demonstrate that both h-TREX1 and m-TREX1 degrade the unprocessed HIV-1 U3 DNA at a faster rate and at higher efficiency than the processed viral DNA substrates.

**Figure 3.**
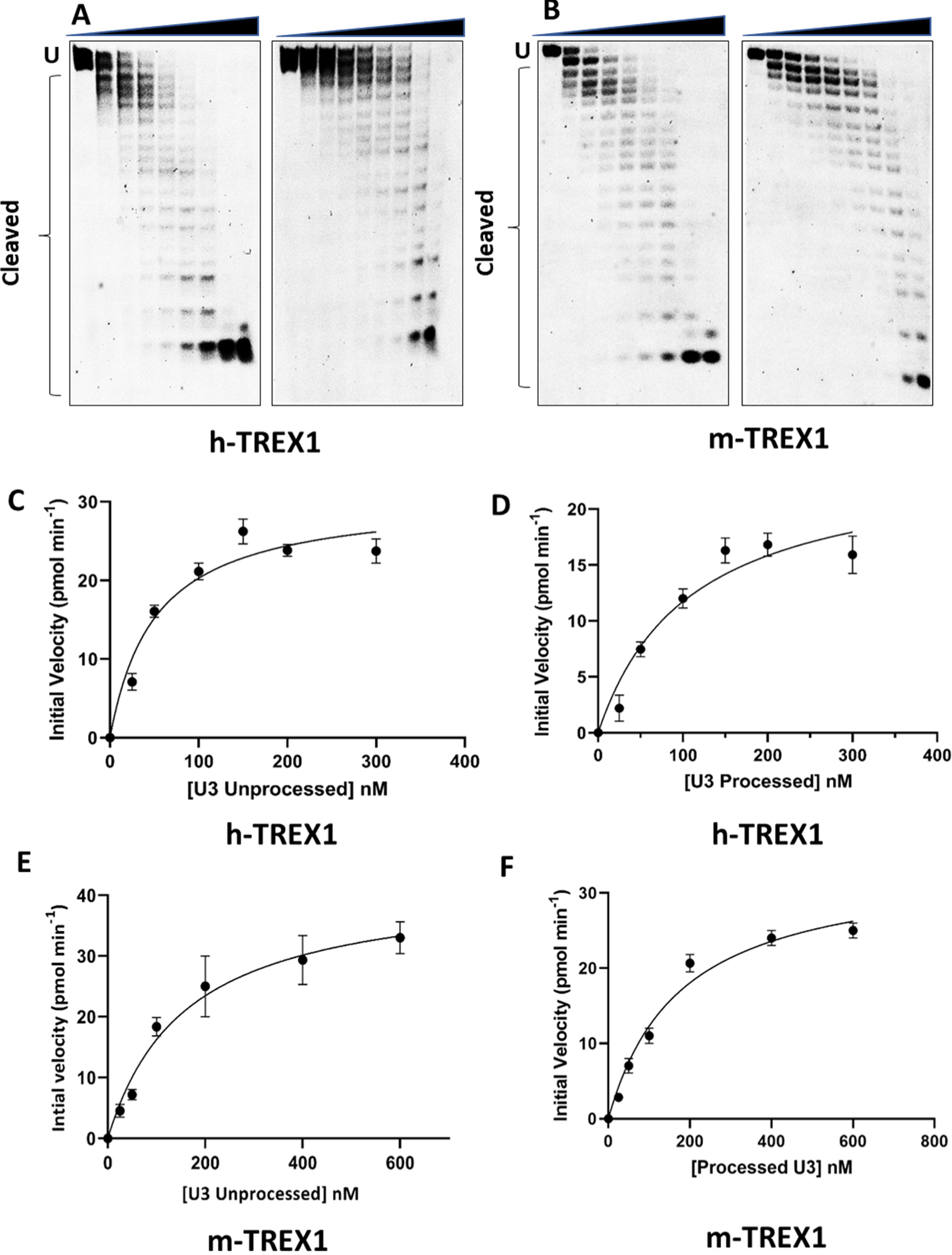
TREX1-mediated degradation of U3 viral DNA substrates. **(A-B)** Representative sequencing gel images showing degradation of unprocessed (left panel) and 3’-processed (right panel) U3 HIV-1 DNA substrates by **(A)** h-TREX1 and **(B)** m-TREX1. Uncleaved (U) and cleaved substrates are highlighted in these images. The uncleaved and cleaved products of both unprocessed and processed viral DNA substrates respectively in panels A and B were separately analyzed and aligned at the same level for the purpose of comparison. The progression curve of the **(C-D)** h-TREX1 and **(E-F)** m-TREX1 enzymatic activity with the U3 unprocessed and U5 3’-processed HIV-1 DNA substrates. Each result and the data points on the kinetic progression curves are the representative of three experimental replicates with errors bars representing SEM.

**Figure 4.**
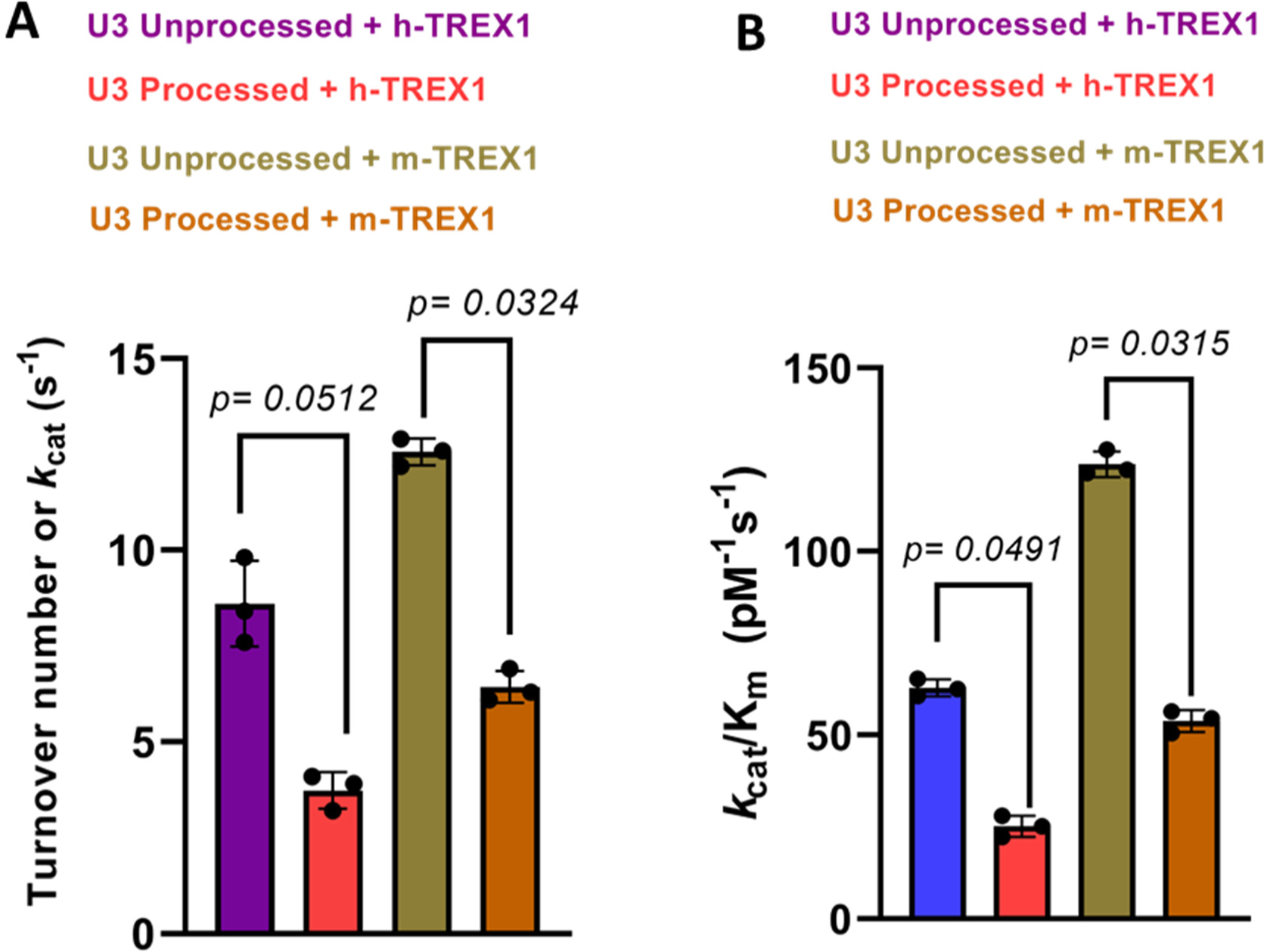
Comparison of the rate of TREX1-mediated degradation of U3 DNA substrates. The bar graphs show the quantitative comparison of the catalytic turnover rate (*k*_cat_) of h-TREX1 and m-TREX1 activity with U3 **(A)** unprocessed and **(B)** 3’-processed HIV-1 DNA substrates. SEM values + /− from three independent experiments are shown. The p-values are shown just above the bar graphs show statistical significance.

**Table 2.**

| | DNA Substrates | $V_{\max}$ (pmol min<sup>-1</sup>) | $K_m$ (nM) | $k_{\text{cat}}$ (sec<sup>-1</sup>)* | $k_{\text{cat}} / K_m$ (pM<sup>-1</sup> s<sup>-1</sup>) |
| --- | --- | --- | --- | --- | --- |
| h-TREX1 | U3 Unprocessed | 39.72 ± 7.5 | 150.71 ± 12.3 | 9.8 ± 0.8 | 65.3 ± 0.45 |
|  | U3 Processed | 25.2 ± 5.1 | 160.4 ± 14.2 | 4.5 ± 0.6 | 28.05 ± 0.71 |
| m-TREX1 | U3 Unprocessed | 31.3 ± 3.1 | 98.7 ± 6.2 | 12.6 ± 0.51 | 127.8 ± 0.61 |
|  | U3 Processed | 20.5 ± 5.1 | 125.3 ± 4.1 | 6.3 ± 0.42 | 54.6 ± 0.65 |

### TREX1 binds to the unprocessed HIV-1 DNA ends with higher affinity compared to the 3’-processed DNA

The turnover rate (*k*_cat_) of a substrate is dependent on the K_M_, which is the substrate concentration required for half-maximal velocity of enzyme activity. However, K_M_ does not provide details about the binding affinity between the substrate and enzyme. Equilibrium dissociation constant (K_d_) measures the rate at which the enzyme-substrate complex is formed and is another critical kinetic parameter in product formation (54). Therefore, we calculated K_d_ values of TREX1 to HIV-1 DNA substrates by Fluorescence polarization (FP) anisotropy (55–58). FP anisotropy is a sensitive assay to measure the binding affinity of ligands to proteins using a fluorophore (Fig. 5A) (59–61). When a fluorophore-labeled ligand binds to a protein, the rotational motion of the protein-ligand complex is reduced compared to the free ligand (Fig. 5A), resulting in higher anisotropy (61–63). Exploiting this unique property of FP anisotropy, we measured the binding affinity of TREX1 to the unprocessed and 3’-processed substrates. For this assay, high protein concentration is required, that could only be achieved with our m-TREX1 preparation. Therefore, we titrated a fixed concentration of each of the viral DNA substrate with increasing concentration of m-TREX1. The change in anisotropy of the unprocessed and 3’-processed substrates were plotted as a function of m-TREX1 concentration (Fig. 5B). The resulting binding curve(s) showed that FP of the unbound substrates increased in a dose-dependent manner as a function of m-TREX1 concentration (Fig.5B). Accordingly, complete binding of labelled substrates to m-TREX1 was indicated by maximal polarization values at the highest protein concentration used (Fig. 5B). Then, the K_d_ value of m-TREX1 for each substrate was calculated by fitting the FP data into a non-linear regression model of one-site specific saturation binding. These analyses revealed that the unprocessed viral DNA substrate (both U5 and U3) has an approximately 2 to 2.5-fold lower K_d_ value indicative of higher binding affinity to m-TREX1 than the 3’-processed viral DNA substrates (Fig. 5C, Table-3). For example, the K_d_ of unprocessed U5 DNA to m-TREX1 was ∼76 nM, compared to ∼179 nM for the 3’-processed viral DNA (Table-3). Similarly, the unprocessed U3 substrate bound to m-TREX1 with a K_d_ of ∼118 nM compared to a value of ∼215 nM for the 3’-processed U3 substrate (Table 3). These results demonstrate that the unprocessed viral DNA substrates from both the ends of HIV-1 LTR form a complex with TREX1 with higher binding affinity over the 3’-processed viral DNA. Interestingly, a comparative analysis of the K_d_ values with the corresponding K_M_ values of m-TREX1 (Tables 1-2) revealed that the K_M_ values of the unprocessed DNA strand were higher than the K_d_ values (i.e., for U3 unprocessed-K_M_ of 150 nM vs K_d_ of 118 nM and for U5 unprocessed-K_M_ of 109 nM vs K_d_ of 76 nM). Conversely, for the 3’-processed strand the K_M_ values were lower than the K_d_ values (i.e., for U3 processed-K_M_ of 160 nM vs K_d_ of 215 nM and for U5 processed-K_M_ of 90 nM vs K_d_ of 179 nM). Importantly, a relative higher ratio of K_M_/K_d_ (Table 4) is an indication that the equilibrium of the enzyme-substrate complex favors higher rate of product formation with tighter binding, whereas a lower ratio is indicative of a slower rate of product formation (64). These results suggest favorable kinetic and thermodynamic properties that contribute to the preferential degradation of the unprocessed HIV-1 DNA by TREX1.

**Figure 5.**
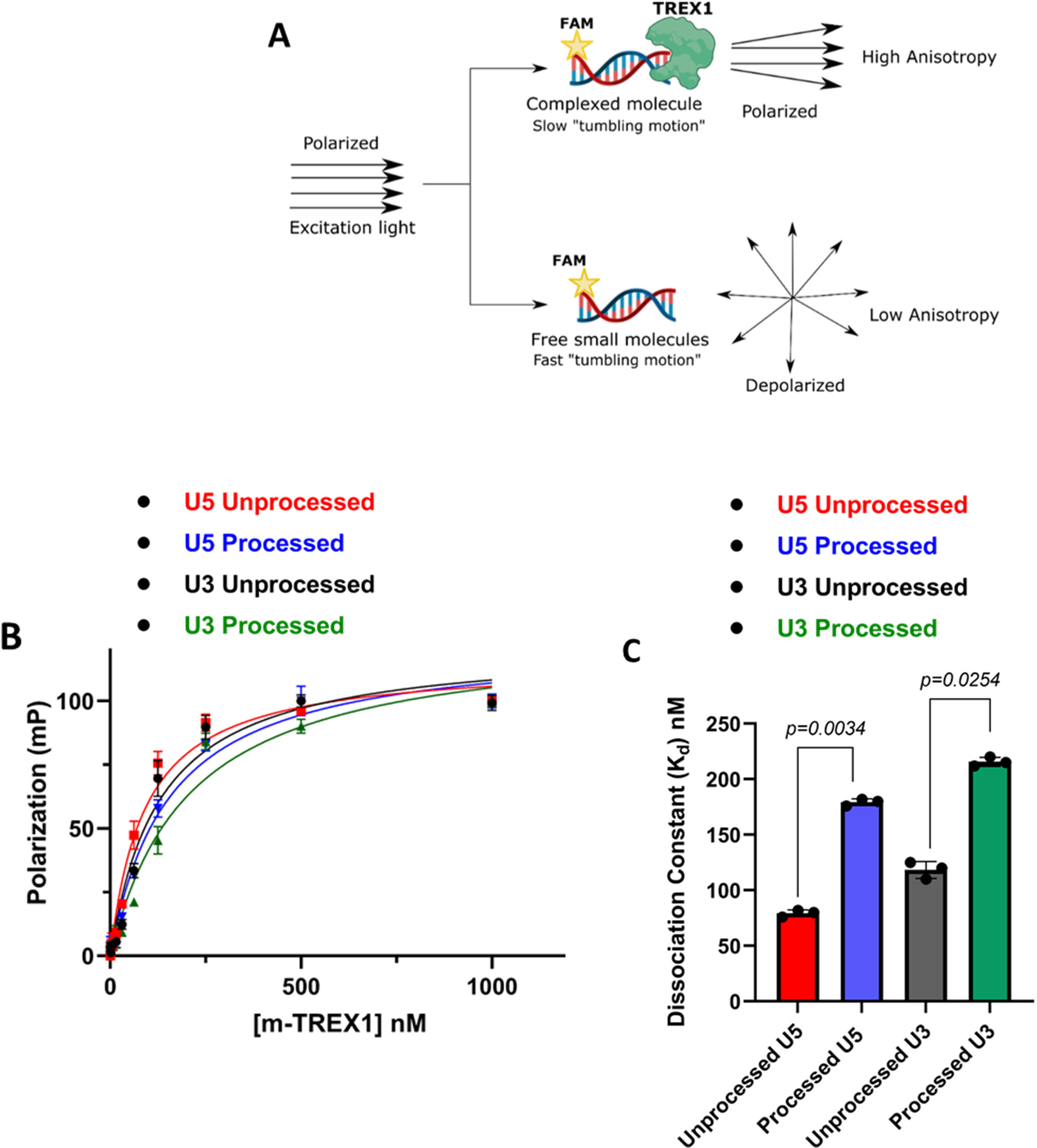
Measurement of HIV-1 DNA binding affinity to TREX1 using fluorescence polarization anisotropy. **(A)** Schematic representation of principle Fluorescence Anisotropy used in this study. **(B)** Comparison of saturation binding curve of m-TREX1 complexed with unprocessed and processed HIV-1 DNA substrates. **(C)** Bar graph representation of binding affinity (dissociation constant) values of unprocessed and processed HIV-1 DNA substrate complexed with TREX1. The K_d_ data shown here was obtained from three independent experimental replicates with error bars representing the SEM.

**Table 3:** Dissociation constant of m-TREXl.

| Viral DNA substrate | U5 Unprocessed | U5 Processed | U3 Unprocessed | U3 Processed |
| --- | --- | --- | --- | --- |
| Dissociation constant ($K_d$) nM | 76.3 ± 7.5 | 179.3 ± 6.5 | 118.4 ± 8.4 | 215.6 ± 4.6 |

**Table 4:** K_m_/K_d_ ratio of m-TREXl.

| Viral DNA substrate | U5 Unprocessed | U5 Processed | U3 Unprocessed | U3 Processed |
| --- | --- | --- | --- | --- |
| $K_m/K_d$ ratio | 1.5 ± 0.02 | 0.52 ± 0.01 | 0.49 ± 0.02 | 0.28 ± 0.05 |

### The unprocessed HIV-1 DNA is proximally and favorably positioned in the active site of TREX1

To further probe the details of the selective degradation of the unprocessed HIV-1 DNA ends by TREX1, we analyzed the structural interaction between the active site residues of the enzyme and the substrates. Particularly, to pin-point the molecular interactions between TREX1 and the unprocessed and 3’-processed HIV-1 DNA ends, we carried out *in silico* studies using the recent crystal structure of h-TREX1 (PDB ID: 7TQQ) and the refined structure of m-TREX1 (PDB ID: 3MXJ). We employed HDOCK docking program that uses Fast Fourier Transform (FFT)-based global sampling of putative binding mode to predict macromolecular interactions (37). HDOCK generated approximately one hundred theoretical models of complexes between h-TREX1 and each of the HIV-1 DNA substrates. These complexes were then ranked based on their docking energies. Scoring function is used for h-TREX1-viral DNA complexes where one binding mode corresponding to the best scored-translation is retained for each rotation. Based on these criteria, docking of TREX1 with the unprocessed and 3-processed viral DNA substrates of both U5 and U3 ends generated top 10 theoretical models. The calculated energies of the selected model of h-TREX1 complexes with unprocessed U5 DNA ends was −140.9 kcal/mol, while that of the 3’-processed U5 DNA end was −125.4 kcal/mol (Table 5). Likewise, the docking energies of the h-TREX1 complexes with the U3 unprocessed and 3’-processed viral DNA were −135.3 kcal/mol and −118.5 kcal/mol, respectively (Table 5). The best docked models of all the TREX1-viral DNA complexes, including those formed by the active site residues were carefully analyzed (Fig. 6). Interestingly, the 3’-ends of the unprocessed viral DNA substrates from both the U5 (Fig. 6A) and U3 ends (Fig. 6C) were positioned in the close conformational proximity to the catalytic residues Asp18, Glu20, Asp200 and His195 of h-TREX1 active site. In contrast, the recessed 3’-end of the processed DNA substrates were oriented in a distal conformation from the catalytic residues (Fig. 6B and 6D). Further, the backbone of the unprocessed DNA strands was considerably stabilized by polar interactions with the C-terminal Lys75, Arg128, Ser176, Ser 190, His 195 residues of h-TREX1, whereas no such stabilizing molecular interactions were observed with the backbone of the 3’-processed DNA ends. Since h-TREX1 and m-TREX1 share high sequence and structural identity (28,52,53), we also analyzed the secondary structure similarity (SSM) superimposition of crystal structures of m-TREX1 with h-TREX1 complexed with U5 unprocessed (Fig. 6E), U5 3’-processed (Fig 6F), U3 unprocessed (Fig. 6G) and U3 3’-processed (Fig. 6H). The SSM results revealed a low RMSD value of <1.0 (0.75 Å), suggesting no significant differences in the Cα-backbones between both the TREX1. Also, we observed similar mode of binding of m-TREX1 with the substrates, where the unprocessed viral DNA is oriented in proximity of the active site residues, whereas the 3’-processed viral DNA is positioned distally. These results illustrate that TREX1 active site retains favorable conformation with the unprocessed viral DNA to mediate efficient catalysis, while a distally oriented 3’-processed HIV-1 DNA is sterically less accessible to the catalytic residues. These molecular interactions capture novel structural and conformational details underlying the slower rate of degradation of the 3’-processed substrates by TREX1.

**Figure 6.**
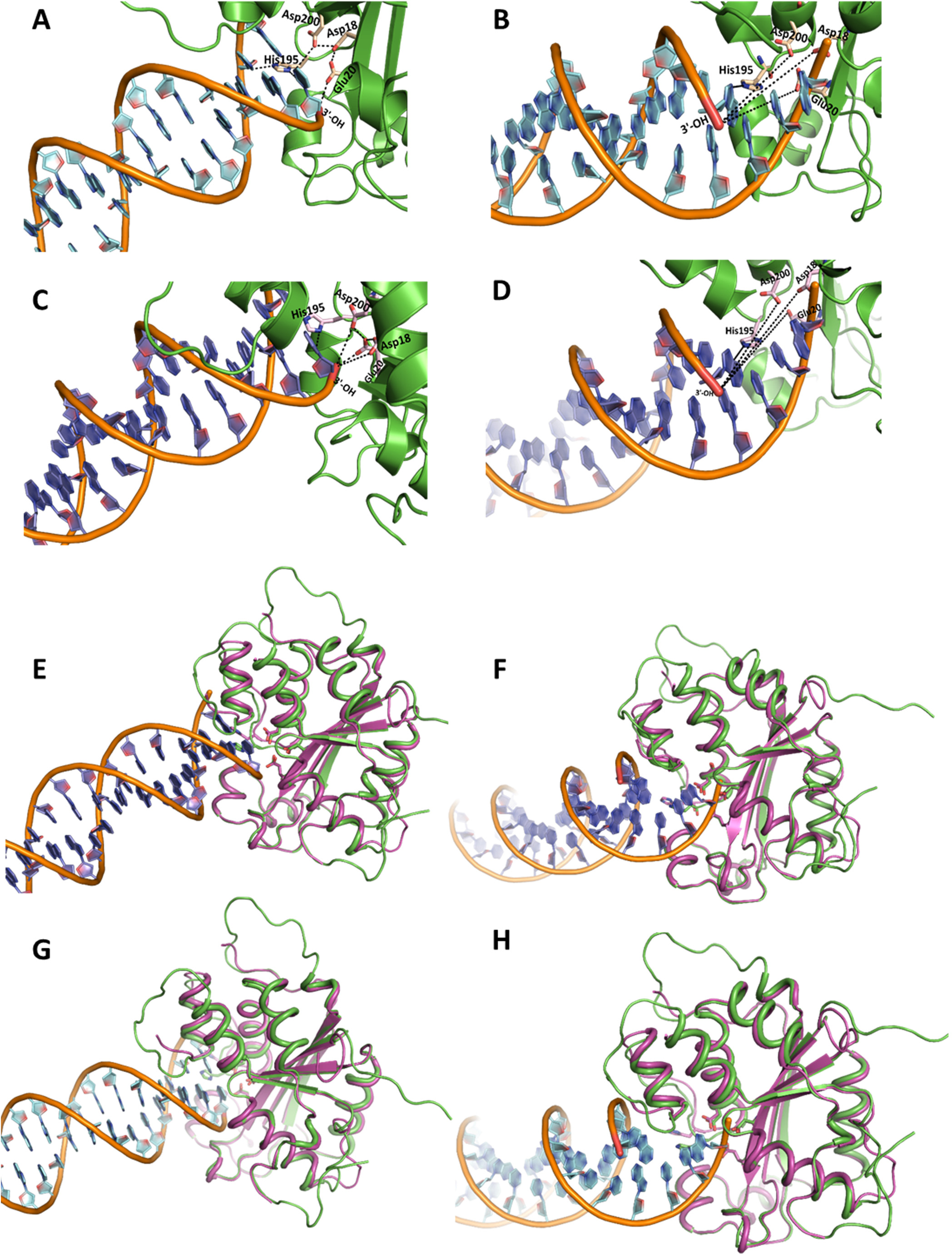
Structural interactions of viral DNA substrates with TREX1 active site. Cartoon representation of (**A**) U5 unprocessed viral DNA and (**B**) U5 3’-processed viral DNA binding with h-TREX1 active site. Nucleotide base rings are shown in cyan, DNA helices with brown backbone, h-TREX1 (PDB ID: 7TQQ) active site region is in green ribbons, the 3’-OH ends of the DNA are in red, and the active site residues are shown in light yellow sticks. The 3’-end (red) of the unprocessed DNA forms proximal contacts while 3’-OH of the processed DNA is distantly oriented from the h-TREX1 active site residues (Asp18, Asp200, Glu20 and His195 labeled in black). The black dashed lines indicate polar proximities of 3’-end of U5 viral DNA from the constellation of active site residues of h-TREX1. **(C)** U3 unprocessed- and **(D)** U3 3’-processed viral DNA binding mode with h-TREX1 active site. Nucleotide base rings are shown in purple, DNA helices with brown colored backbone, h-TREX1 active site region is in green ribbons, 3’-OH ends in red and the active site residues are in light magenta sticks. The binding mode of the active site of h-TREX1 illustrate that 3’-end of the unprocessed DNA forms a similar proximal contact as seen with U5 unprocessed substrate, while 3’-OH end of the processed DNA distantly oriented from the h-TREX1 key residues of active site. The black dashed lines denote the distancing of 3’-OH end from the active site residues. **(E-H)** The structural superposition showing the highly conserved overall secondary structure folds of both m-TREX1 and h-TREX1. (**E**) Structural superpositions of the m-TREX1 shown in magenta (PDB ID: 3MXJ) with the h-TREX1 (green cartoon) complexed with U5 unprocessed, (**F**) U5 processed, (**G)** U3 Unprocessed and (**H**) U3 processed viral DNA ends. All protein-DNA docking was performed using HDOCK.

**Table 5:**
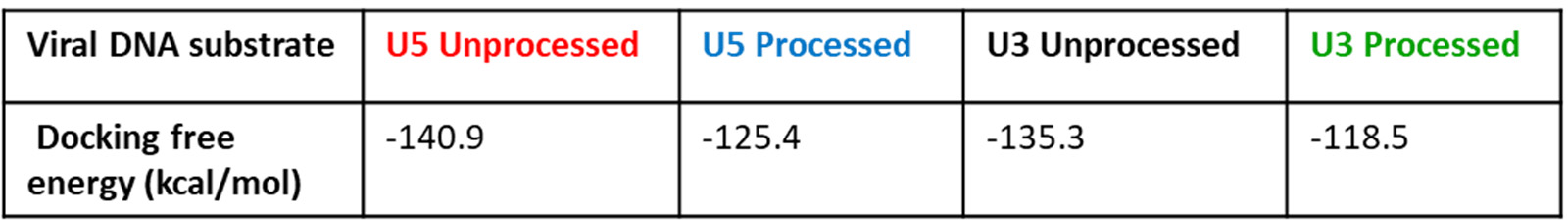
Docking free energy values of h-TREXl complexed with unprocessed and processed viral DNA.

| Viral DNA substrate | U5 Unprocessed | U5 Processed | U3 Unprocessed | U3 Processed |
| --- | --- | --- | --- | --- |
| Docking free energy (kcal/mol) | -140.9 | -125.4 | -135.3 | -118.5 |

### TREX1 forms stable complex with the unprocessed HIV-1 DNA compared to the 3’-processed HIV-1 DNA

Our docking results showed distinct molecular interactions between the unprocessed and processed HIV-1 DNA with TREX1 active site residues (Fig. 6). To further understand the binding mode and stability of h-TREX1 with these substrates, we carried out molecular dynamics simulation studies using Gromacs2021.5 that consider charges of the biomolecules to study macromolecular interactions as a function of time (43). First, we calculated the root mean square deviation (RMSD) of the protein backbone atoms to examine the stability of the TREX1-viral DNA complexes (Fig. 7A). The RMSD plot revealed that all the systems are well equilibrated well before ∼ 200 ns of simulation. These simulations demonstrated that TREX1 in its apo-form has a higher RMSD value (0.7-0.8 nm) after the equilibration time suggestive of the process of sampling of the apo-enzyme. Interestingly, TREX1 complexed with U3 and U5 unprocessed DNA were highly stable with a lower RMSD value of 0.5-0.6 nm (Fig. 7A). In contrast, TREX1 complexed with the 3’-processed DNA retained higher RMSD with fluctuations between ∼ 0.6 to 0.8 nm (Fig. 7A). Notably, the complex of U5 3’-processed strand retained the higher level of fluctuation immediately after the equilibrium time (∼200 ns). However, the complex of the U3 3’-processed strand fluctuated below 0.6 nm for a longer period before attaining maximum fluctuation by ∼400 ns, suggesting distinct conformational modes of this complex.

**Figure 7.**
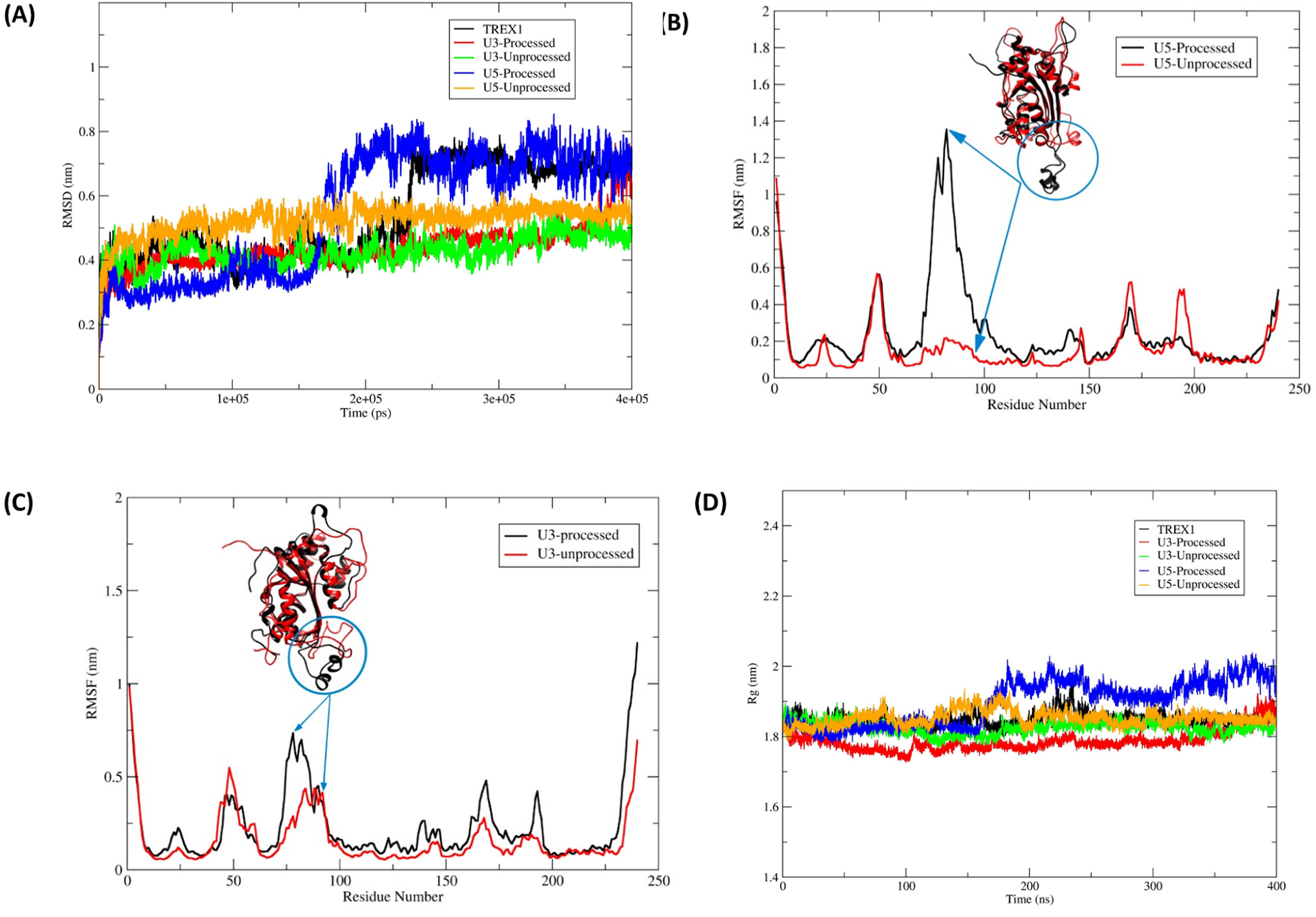
Molecular dynamics simulation of TREX1 complexed with HIV-1 DNA substrates. **(A)** RMSD for backbone atoms of the h-TREX1 during molecular dynamics simulations for 400 ns. **(B-C)** RMSF of the backbone Cα atoms of h-TREX1 with **(B)** U5 and **(C)** U3 HIV-1 DNA substrates during molecular dynamics simulations for 400 ns. The highly flexible regions show higher RMSF value (Region; 70 to 90) while the constrained regions show low RMSF value. The RMSF value of h-TREX1 complexed with **(B)** U5 and **(C)** U3 processed DNA is higher as compared to the complex with **(B)** U5 and **(C)** U3 unprocessed DNA complexes. **(D)** Rg of h-TREX1 during molecular dynamics simulations for 400 ns time step. The Rg value of TREX1 complexed with U3 processed and U5 processed DNA is higher compared to the TREX1 complex with U3 and U5 unprocessed DNA.

To further study these distinct structural changes, we calculated the root mean square deviation (RMSF) and the radius of gyration (Rg) in the TREX1 and HIV-1 DNA complexes (Fig. 7B-D). The RMSF plots revealed that binding of the 3’-processed DNA ends in the TREX1 catalytic region attained a highly flexible conformation indicative of an unstable complex (Fig. 7B and Suppl. movies 1 and 3). In contrast, the conformation of the catalytic region was highly constrained and stable with the unprocessed DNA (Fig. 7C and Suppl. movies 2 and 4). Then, the compactness of structural changes in TREX1 was studied using the Rg values. Interestingly, TREX1 complexed with the unprocessed DNA ends showed a lower Rg value between 1.8 nm to 1.85 nm, whereas with the 3’-processed DNA a higher Rg value between 1.9 nm to 2.0 nm was obtained (Fig. 7D). These results suggest that TREX1 is in a compact state with the U5 unprocessed DNA compared to the respective 3’-processed DNA. Interestingly, it becomes evident that at early time points of the simulation, the Rg value of the 3’-processed U3 DNA is relatively lower when compared to the U3 unprocessed DNA. However, after 350 ns, the Rg value of the 3’-processed U3 DNA starts to sharply increase and achieves a value indicative of an unstable complex. These results indicate that the complex of TREX1-unprocessed substrate is highly stable, while the complex with the 3’-processed DNA is unstable. Taken together, these analyses highlight the structural details that may facilitate preferential degradation of a subset of HIV-1 DNA substrates by TREX1.

## Discussion

TREX1 degrades endogenous cellular DNA to prevent innate immune response to self-DNA (50,65,66). TREX1 activity is also usurped by HIV-1 as a mechanism to evade the cellular antiviral ISD response. For instance, synthesis of HIV-1 DNA by reverse transcription fails to trigger ISD response in an infected cell (67). Yet, in the absence of TREX1, HIV DNA accumulates and induces robust ISD response (68–70). Additionally, knockdown of TREX1 inhibits HIV-1 infection, suggesting a critical role of this cellular exonuclease in HIV-1 life cycle (71). It is predicted that HIV-1 DNA levels are maintained below a threshold level by TREX1 to avert detection by the cellular sensors. Interestingly, TREX1 activity has to be precisely regulated to prevent complete degradation of HIV-1 DNA, since viral DNA integration into the host genome is required for HIV-1 infection. However, the mechanism by which a subset of HIV-1 DNA that are integration-competent is protected from TREX1 activity remains largely unknown. Nevertheless, there is evidence that viral DNA could be protected by the binding of the PIC-associated IN (72). The viral capsid could also protect the viral DNA as recent studies suggest that reverse transcription is completed within an operationally intact capsid (73). Importantly, we recently discovered that the integration-competent HIV-1 DNA alone (without IN) remains resistant to TREX1-mediated degradation, whereas the integration-incompetent viral DNA is efficiently degraded (27). Here, we describe the kinetic and structural mechanisms of the preferential degradation of the integration-incompetent HIV-1 DNA ends and uncover the molecular features of the integration-competent HIV-1 DNA those confer resistance to TREX1 activity.

HIV-1 integration relies on the sequential steps of 3’-processing of the viral DNA ends followed by the strand transfer (20). The 3’-processed viral DNA with a recessed 3’-CA_OH_ end is competent for the strand transfer step of integration, whereas the unprocessed blunt-ended viral DNA is integration-incompetent (20). Remarkably, our exonuclease activity results revealed that the unprocessed HIV-1 DNA of both U5 and U3 ends of the LTR are preferentially degraded by TREX1 when compared to the 3’-processed viral DNA substrates (Fig. 1-4). We have previously described that the presence of the recessed 3’-ends with a 2nt 5’-overhang of the processed HIV-1 DNA is not responsible for this preferential degradation, since DNA substrates with either a blunt-end or a 3’-recessed end were efficiently degraded by TREX1 (27). Since TREX1 is known to degrade a variety of DNA substrates (3,49,74,75), the resistance of 3’-processed HIV-1 DNA to degradation was unexpected. Nevertheless, TREX1 activity could be influenced by the secondary structure of the substrate (75,76) and sequence preference (77). Therefore, we probed whether the 3’-processed (integration-competent) and unprocessed (integration-incompetent) HIV-1 DNA possess unique biochemical and/or structural features to alter TREX1 activity. To test this, we combined kinetic studies of exonuclease activity with structural analysis of the complexes of TREX1 with unprocessed and 3’-processed HIV-1 DNA substrates.

Our kinetic analysis revealed that the blunt-ended unprocessed HIV-1 DNA from both the U5 and U3 ends were efficiently degraded, whereas the 3’-processed DNA remained resistant to TREX1 activity (Figs. 1-4). For example, the turnover rate (*k*_cat_) and the catalytic efficiency (*k*_cat_/K_M_) of the unprocessed HIV-1 DNA were significantly higher compared to the 3’-processed DNA by both h-TREX1 and m-TREX1 (Table-1-2). Similarly, the maximum velocity (V_max_) of TREX1 activity was higher for the unprocessed substrates relative to the 3’-processed substrates. Interestingly, the K_M_ value of the U5 unprocessed DNA was higher compared to the 3’-processed strands for both the enzymes (Table-1-2). K_M_ is the substrate concentration at which the initial catalytic rate is half of the maximum velocity of the enzyme activity. Although K_M_ can contribute to the turnover rate, binding affinity between the substrate and enzyme also plays a critical role in catalysis. Accordingly, we observed that the unprocessed viral DNA substrates (both U5 and U3) bind to TREX1 at higher affinity compared to the 3’-processed viral DNA substrates (Table-3). Importantly, a higher ratio of K_M_/K_d_ (Table 4) also provided further evidence that the equilibrium of the TREX1-unprocessed substrate complex favored faster rate of product formation with tighter binding, whereas a lower ratio of K_M_/K_d_ for the TREX1-processed substrate complex was indicative of a slower rate of product formation (64). We are cognizant that the K_d_and K_M_/K_d_ values were calculated for the m-TREX1 and these values for the h-TREX1 could be similar or divergent. Still, these results strongly suggest favorable kinetic and thermodynamic properties that contribute to the preferential degradation of the unprocessed HIV-1 DNA by TREX1.

The higher efficiency of cleavage of the unprocessed HIV-1 DNA substrates containing either the U5 or the U3 ends of the LTR, also indicated that the sequence specificity minimally affects TREX-1 mediated cleavage rate. Interestingly, a faster degradation of the unprocessed HIV-1 DNA observed with both the human and mouse TREX1 also suggested that the TREX1 activity is biochemically conserved. This is most likely due to the high degree of structural and functional similarity between these two enzymes (28,52,53). However, the turnover rates and catalytic efficiency of the two enzymes (Fig. 4A-B) suggested that h-TREX1 is more efficient in cleaving the unprocessed U3 substrate when compared to m-TREX1 (Fig. 4A-B). This observation suggested a degree of species level biochemical difference between the two TREX1 enzymes. Despite of this minor difference, the 3’-processed HIV-1 DNA substrates remained resistant to both the enzymes. Collectively, these kinetic studies clearly demonstrate that both h-TREX1 and m-TREX1 degrade the integration-incompetent unprocessed viral DNA ends at a significantly faster rate than the integration-competent 3’-processed viral DNA.

To gain structural insights into the preferential degradation of a specific subset of HIV-1 DNA, we analyzed the TREX1-DNA complexes by molecular docking and dynamics simulations. Our molecular docking studies identified specific molecular interactions for the preferential activity of TREX1 with unprocessed over 3’-processed HIV-1 DNA (Fig. 6). Specifically, the spatial orientation of unprocessed DNA in the TREX1 active site was proximally located to the active site residues to access the 3’-OH end of the substrate. We predict that this favorable orientation renders TREX1 for optimal catalysis of the unprocessed DNA strands. In contrast, the 3’-processed HIV-1 DNA was distally situated in the binding pocket of TREX1, thereby the 3’-OH end was limited to the exterior region of the active site residues. In this mode of binding, catalysis of 3’-processed DNA is most likely not efficient as the unprocessed HIV-1 DNA. These structural orientations of the substrates in the active site are also conserved between both the h-TREX1 and m-TREX1 as demonstrated by out SSM superimposition studies. These overall conserved interactions most likely explain the similar preference of activity for the HIV-1 DNA substrates. Additionally, our molecular dynamic studies provided further molecular and structural details about the mechanism of preferential activity of TREX1 (Fig. 7). These results illustrated that TREX1 active site retains a favorable conformation when complexed with the unprocessed viral DNA, while a distally oriented 3’-processed HIV-1 DNA is sterically less accessible to the catalytic residues. These distinct structural and conformational interactions may be responsible for the slower rate of degradation of the 3’-processed substrates by TREX1. Our molecular dynamics studies also indicated that the TREX1 complexed with the unprocessed DNA is highly stable, while complex with the 3’-processed DNA is unstable (Fig. 7). Particularly, the unprocessed DNA ends formed a higher number of stable hydrogen bonding interactions with the amino acid residues of TREX1 active site compared to the 3’-processed DNA. Unexpectedly, the complex formation between TREX1 and the 3’-processed DNA of the U3 region required an extended equilibration time compared to the respective U5 substrate. This most likely reflects unique structural features of the U3 DNA ends of HIV-1 LTR that may affect TREX1 binding and activity. Interestingly, these observations are in accordance with the higher K_M_ values of U3 processed substrates compared to the U3 unprocessed substrate (Table 1-2). Crystal structures or CryoEM studies of these complexes are necessary to further pin-point the significance of these unique differences. Nevertheless, these analysis of the TREX1 complexes with the unprocessed and 3’-processed strand reveal the unique molecular interactions that facilitate preferential degradation of a specific subset of HIV-1 DNA substrates.

Based on our results, we propose a hypothetical model for the mechanism by which the integration-competent 3’-processed viral DNA resists TREX1 and is able to integrate into the host genome (Fig. 8). First, in lieu of higher rate of activity, TREX1 efficiently degrades the integration-incompetent unprocessed HIV-1 DNA. In contrast, the 3’-processed (integration-competent) HIV-1 DNA is degraded at a slower rate. The difference in the turn over rate causes a faster clearance of the unprocessed DNA, allows the integration machinery (PIC) to kinetically carry out integration of the 3’-processed HIV-1 DNA into the host genome. It is important to point out that this model does not address how binding of IN to the viral DNA ends may affect the kinetics and interactions between TREX1 and the HIV-1 DNA substrates. Nevertheless, this hypothetical model highlights the kinetic constraints that prevent complete degradation of integration-competent HIV-1 DNA, a mechanism necessary for viral DNA integration and productive HIV-1 infection. Most importantly, this model also explains how TREX1 is utilized during HIV-1 infection to keep the level of total HIV-1 DNA below a threshold level to avoid detection by the cellular sensors to evade the innate immune response.

**Figure 8.**
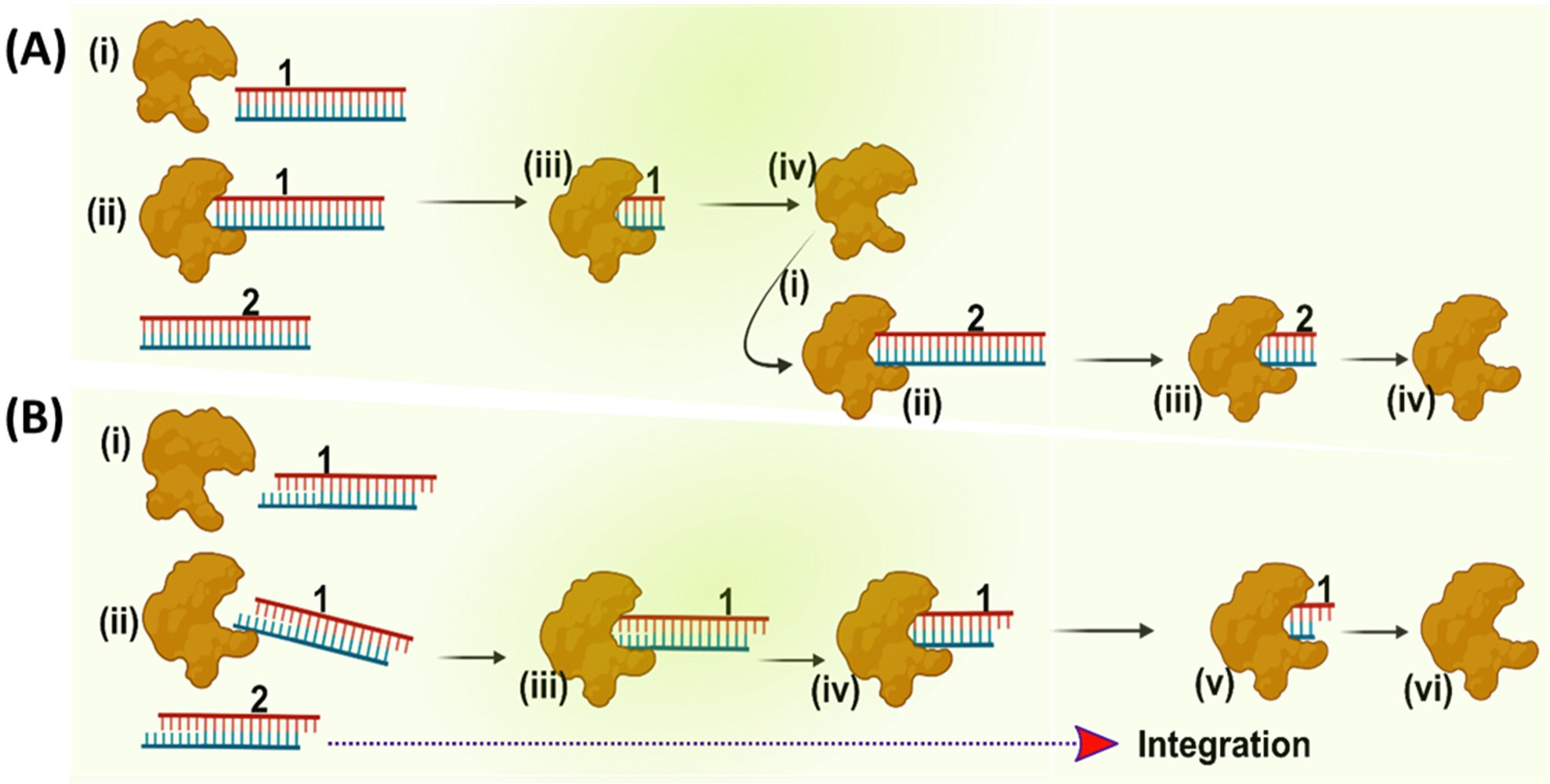
Proposed model of preferential degradation of unprocessed DNA and resistance of 3’-processed DNA to TREX1. **(A)** The panel represents TREX1 activity (generic shape shown in brown) with unprocessed HIV-1 DNA substrate shown in blue-red blunt ended dsDNA. **(B)** Degradation of 3’-processed HIV-1 DNA substrate by TREX1 shown in blue-red sticky ended double strands. A schematic representation showing that TREX1 degrades the unprocessed HIV-1 DNA which is blunt-ended and integration-incompetent at a faster rate, whereas the integration-competent 3’-processed HIV-1 DNA is turned over at a relatively slower rate. This kinetic mechanism allows the 3’-processed HIV-1 DNA to undergo integration into the human genome.

## Data availability

All the data generated in this study are included in the manuscript.

