## Supplementary Figures for "Three Prime Repair Exonuclease 1 preferentially degrades the integration-incompetent HIV-1 DNA through favorable kinetics, thermodynamic, structural and conformational properties"

#### Figure 1: Representative Initial velocity curve analysis of h-TREX1 kinetics with U5 unprocessed and processed HIV-1 substrates

(a)

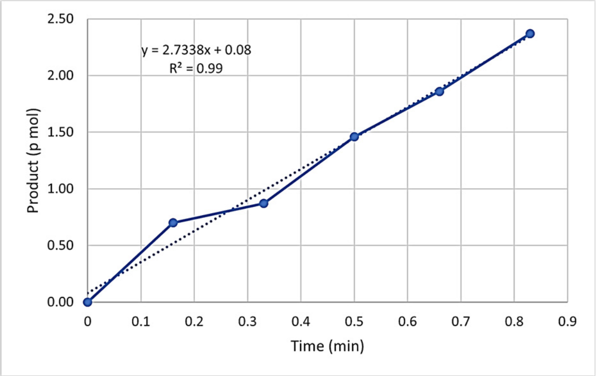


**h-TREX1 with 50 nM U5 unprocessed HIV-1 DNA substrate**

(b)

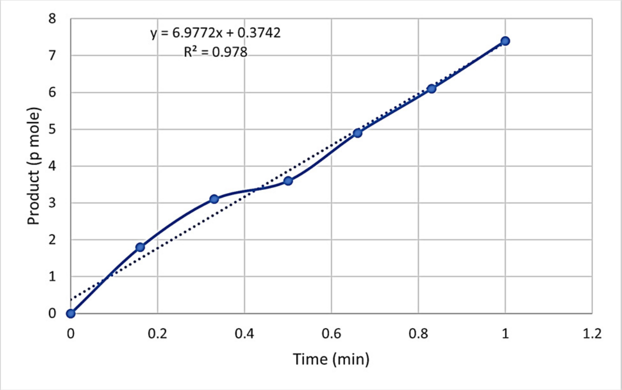


**h-TREX1 with 100 nM U5 unprocessed HIV-1 DNA substrate**

(c)

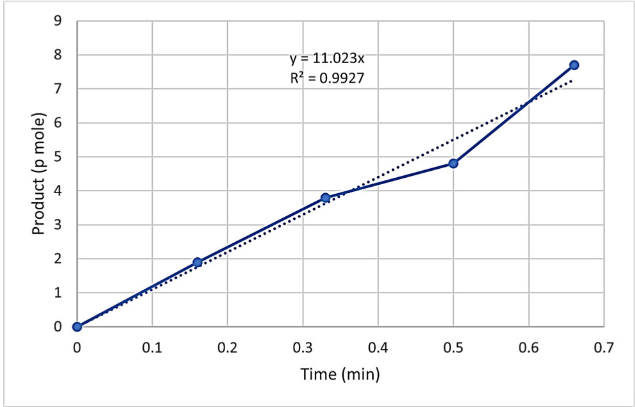


**h-TREX1 with 200 nM U5 unprocessed HIV-1 DNA substrate**

(d)

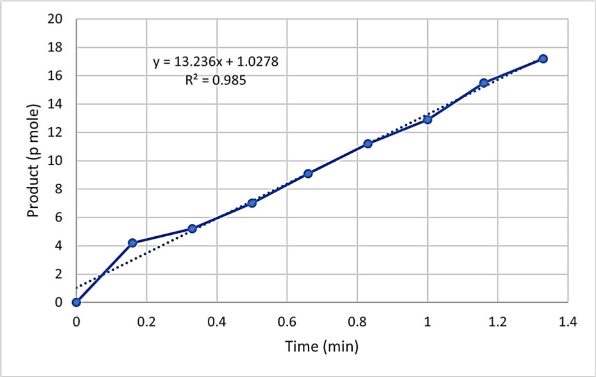


**h-TREX1 with 300 nM U5 unprocessed HIV-1 DNA substrate**

(e)

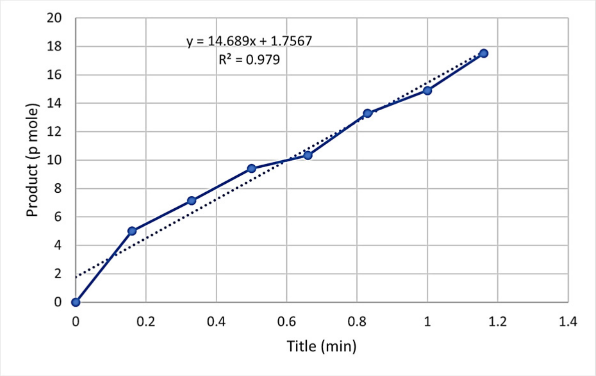


**h-TREX1 with 400 nM U5 unprocessed HIV-1 DNA substrate**

(f)

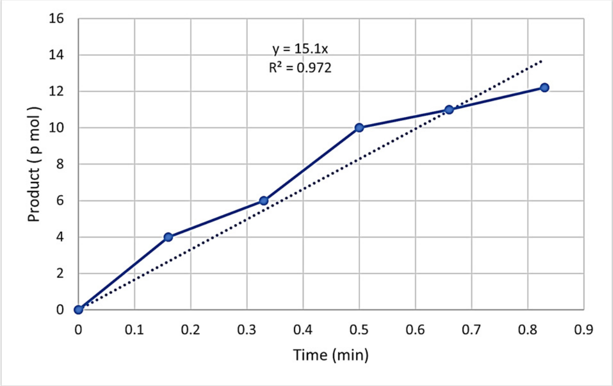


**h-TREX1 with 800 nM U5 unprocessed HIV-1 DNA substrate**

(g)

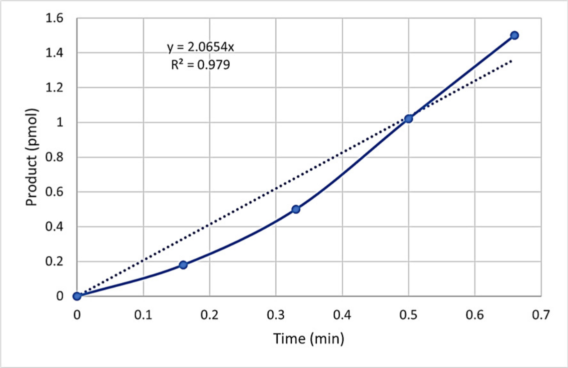


**h-TREX1 with 10 nM U5 Processed HIV-1 DNA substrate**

(h)

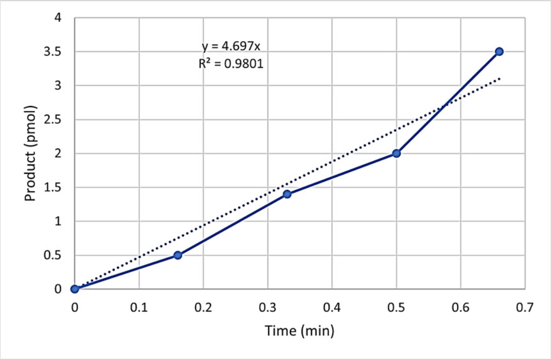


**h-TREX1 with 50 nM U5 Processed HIV-1 DNA substrate**

(i)

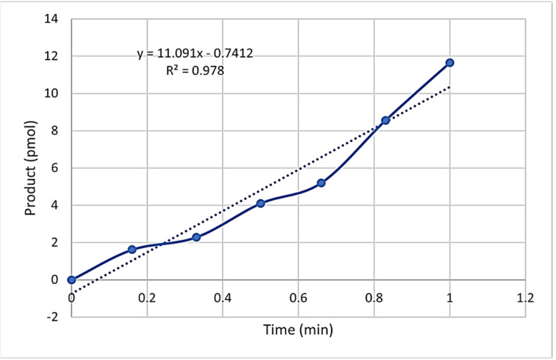


**h-TREX1 with 100 nM U5 Processed HIV-1 DNA substrate**

(j)

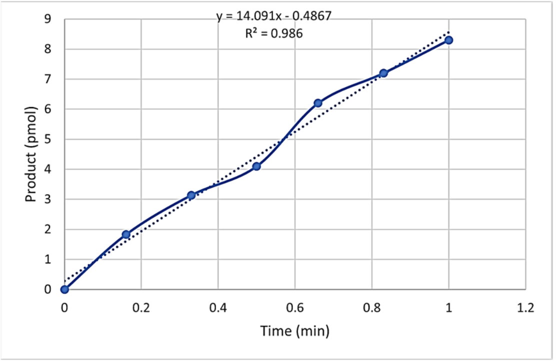


**h-TREX1 with 200 nM U5 Processed HIV-1 DNA substrate**

(k)

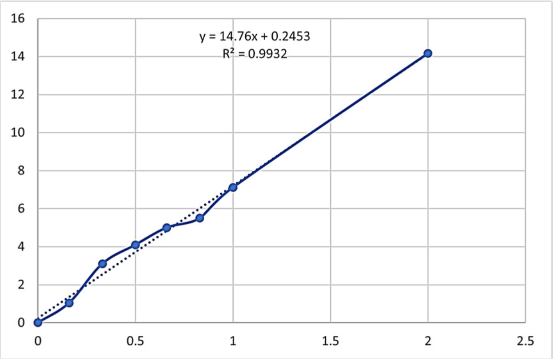


**h-TREX1 with 300 nM U5 Processed HIV-1 DNA substrate**

### Supplementary Figures

#### Figure 2: Representative Initial velocity curve analysis of m-TREX1 kinetics with U5 unprocessed and processed HIV-1 substrates

(a)

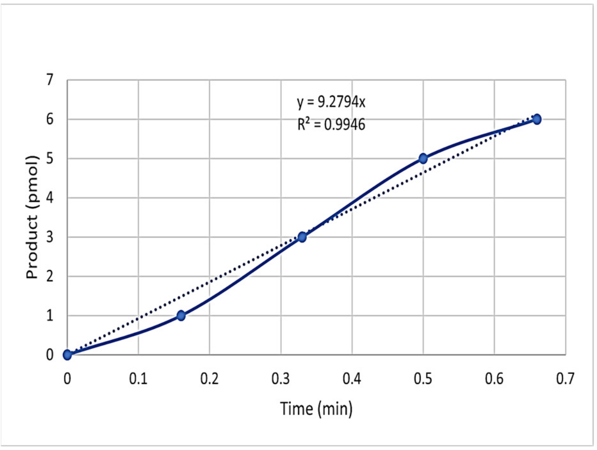


**m-TREX1 with 25 nM U5 Unprocessed HIV-1 DNA substrate**

(b)

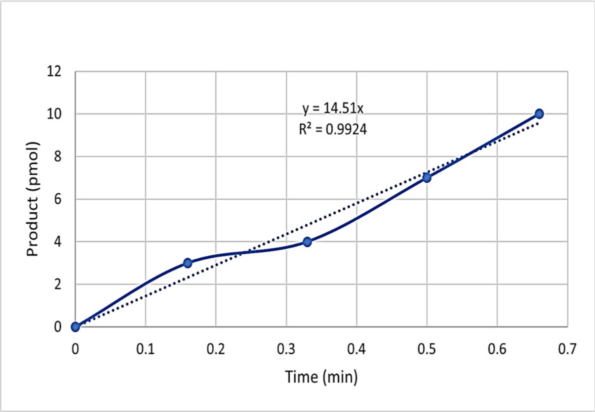


**m-TREX1 with 50 nM U5 Unprocessed HIV-1 DNA substrate**

(c)

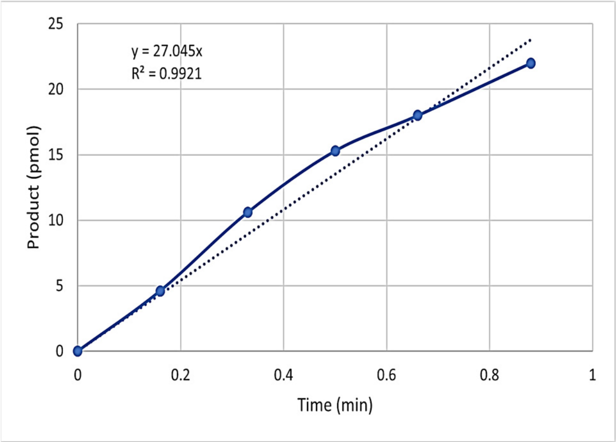


**m-TREX1 with 100 nM U5 Unprocessed HIV-1 DNA substrate**

(d)

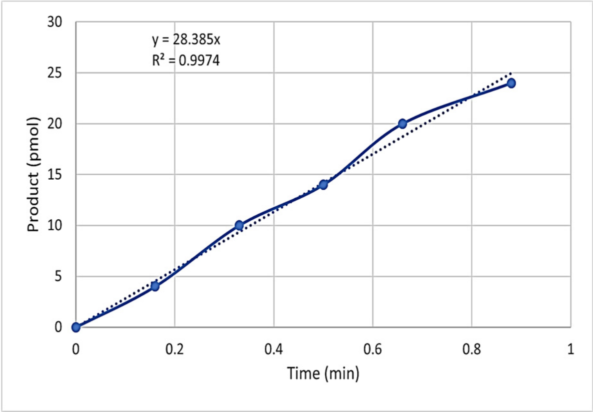


**m-TREX1 with 200 nM U5 Unprocessed HIV-1 DNA substrate**

(e)

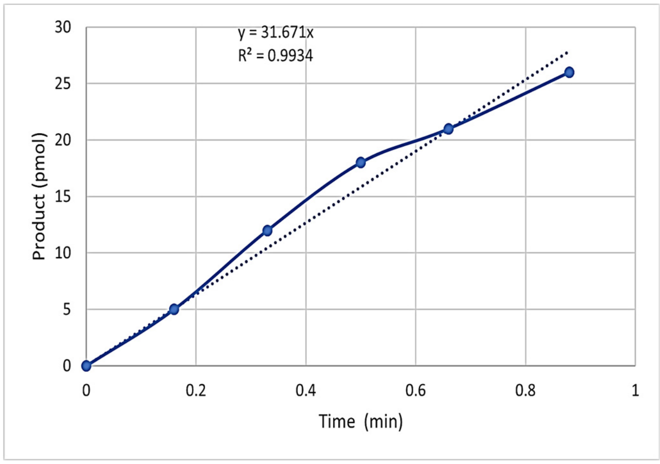


**m-TREX1 with 300 nM U5 Unprocessed HIV-1 DNA substrate**

(f)

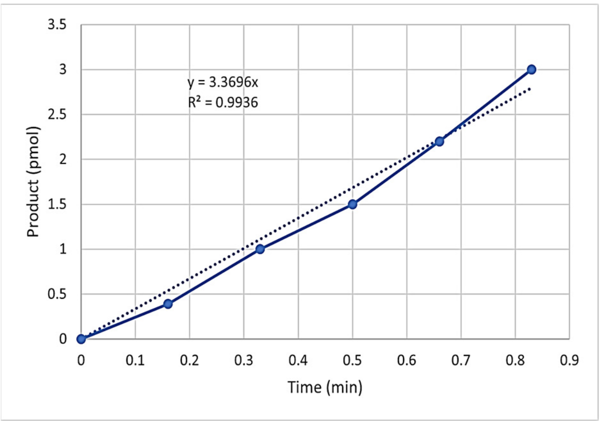


**m-TREX1 with 25 nM U5 Processed HIV-1 DNA substrate**

(h)

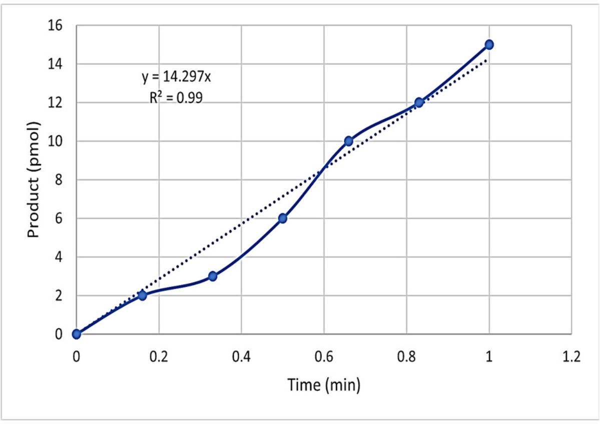


**m-TREX1 with 100 nM U5 Processed HIV-1 DNA substrate**

(g)

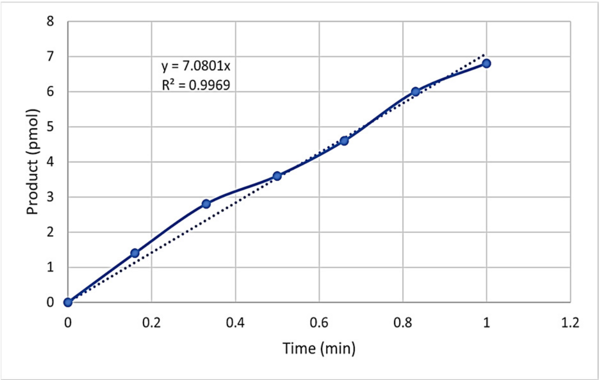


**m-TREX1 with 50 nM U5 Processed HIV-1 DNA substrate**

(i)

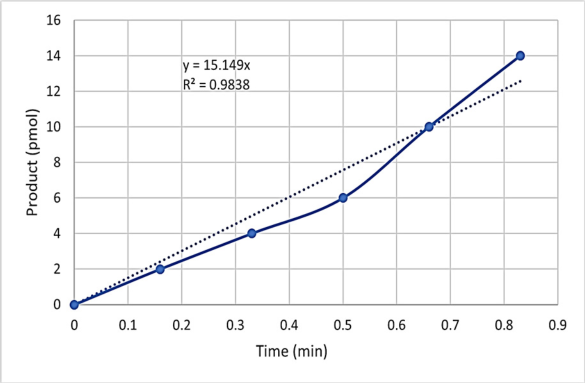


**m-TREX1 with 200 nM U5 Processed HIV-1 DNA substrate**

(j)

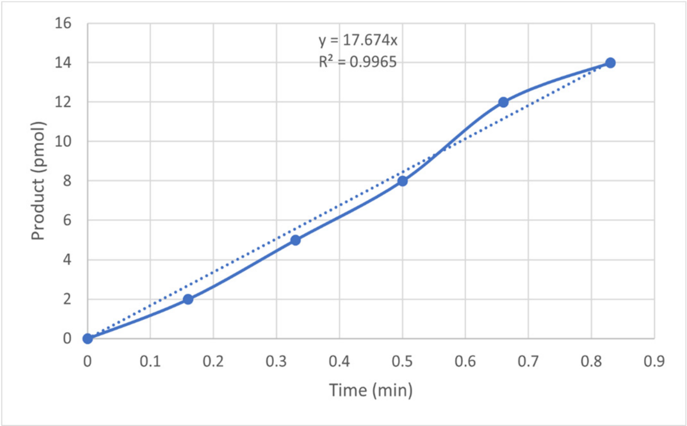


**m-TREX1 with 300 nM U5 Processed HIV-1 DNA substrate**

#### Supplementary Figures

#### Figure 3: Representative Initial velocity curve analysis of h-TREX1 kinetics with U3 unprocessed and processed HIV-1 substrates

(a)

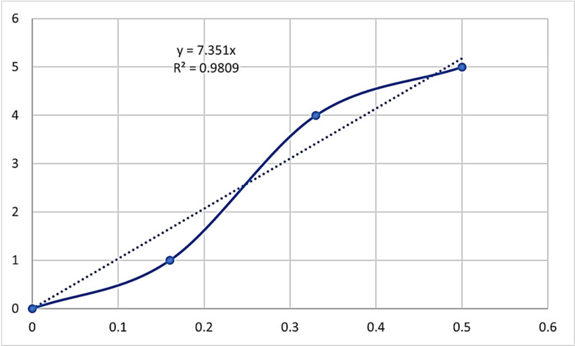


h-TREX1 with 25 nM U3 Unprocessed HIV-1 DNA substrate

(c)

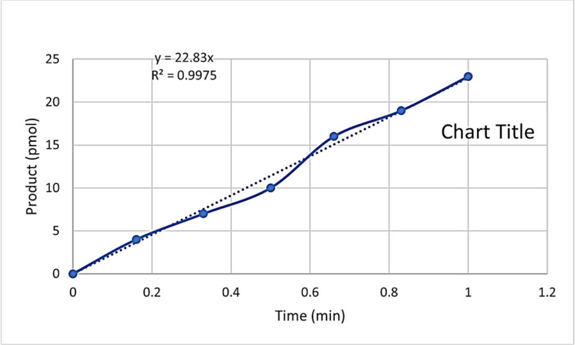


h-TREX1 with 100 nM U3 Unprocessed HIV-1 DNA substrate

(b)

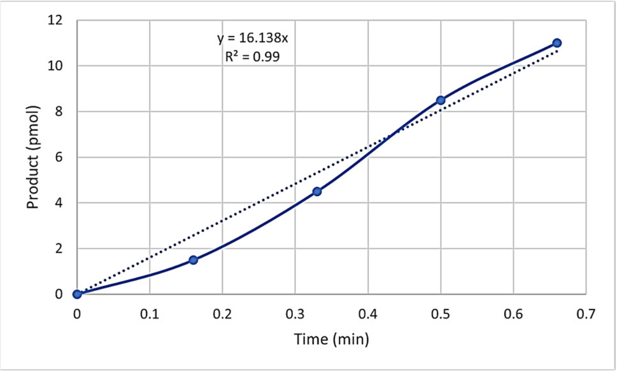


h-TREX1 with 50 nM U3 Unprocessed HIV-1 DNA substrate

(d)

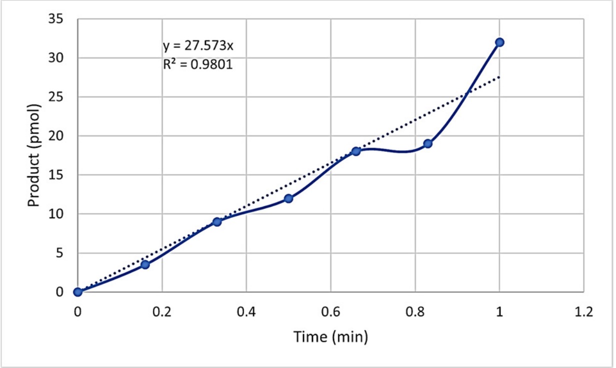


h-TREX1 with 200 nM U3 Unprocessed HIV-1 DNA substrate

(e)

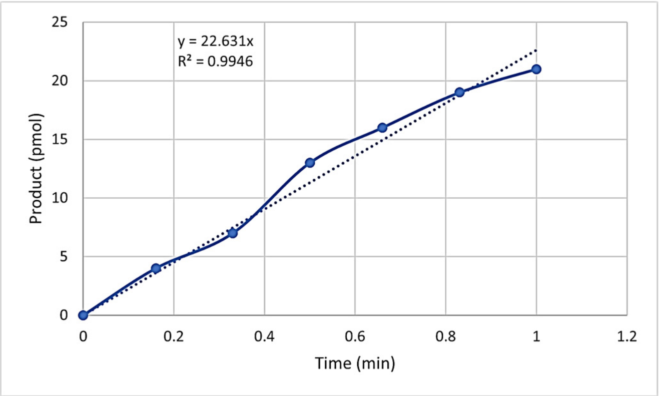


h-TREX1 with 300 nM U3 Unprocessed HIV-1 DNA substrate

**(f)**

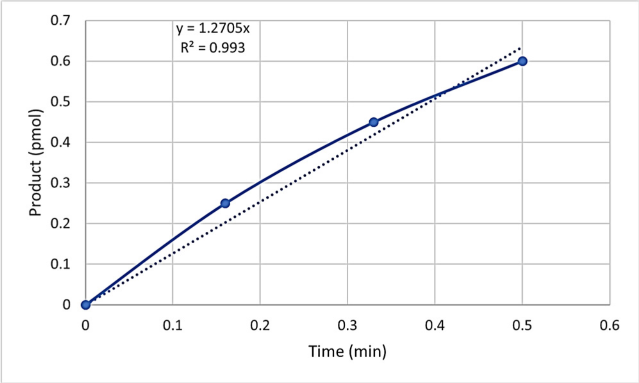


**h-TREX1 with 25 nM U3 Processed HIV-1 DNA substrate**

**(g)**

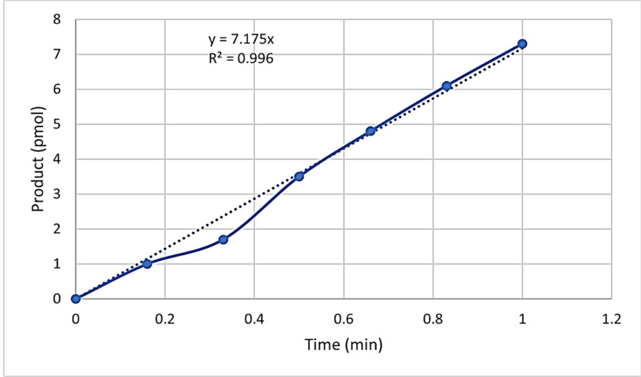


**h-TREX1 with 50 nM U3 Processed HIV-1 DNA substrate**

**(h)**

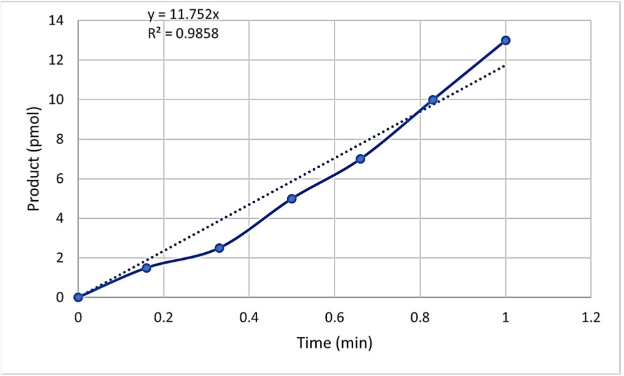


**h-TREX1 with 100 nM U3 Processed HIV-1 DNA substrate**

**(i)**

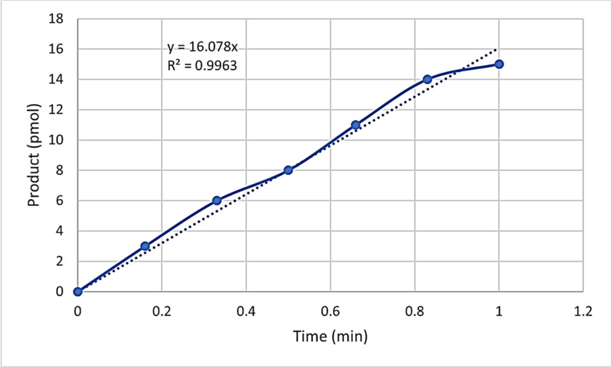


**h-TREX1 with 200 nM U3 Processed HIV-1 DNA substrate**

**(j)**

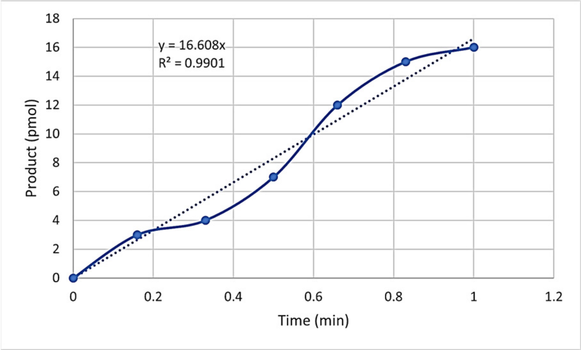


**h-TREX1 with 300 nM U3 Processed HIV-1 DNA substrate**

### Supplementary Figures

#### Figure 4: Representative Initial velocity curve analysis of m-TREX1 kinetics with U3 unprocessed HIV-1 substrates

(a)

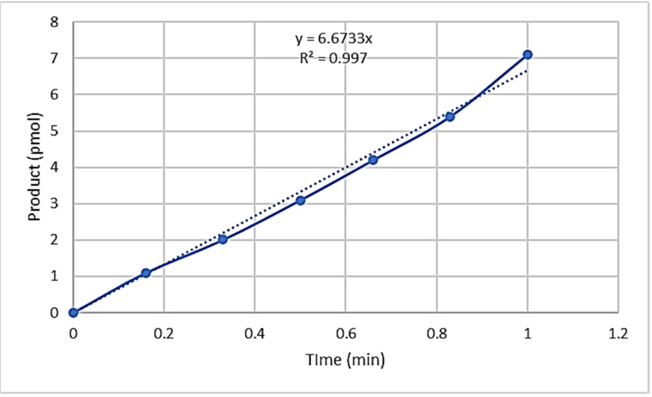


m-TREX1 with 25 nM U3 Unprocessed HIV-1 DNA substrate

(b)

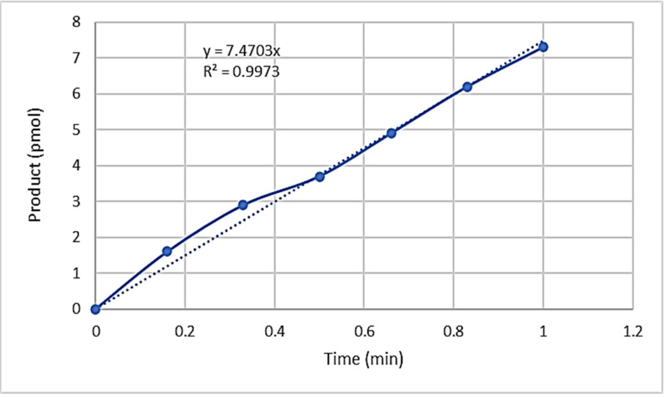


m-TREX1 with 50 nM U3 Unprocessed HIV-1 DNA substrate

(c)

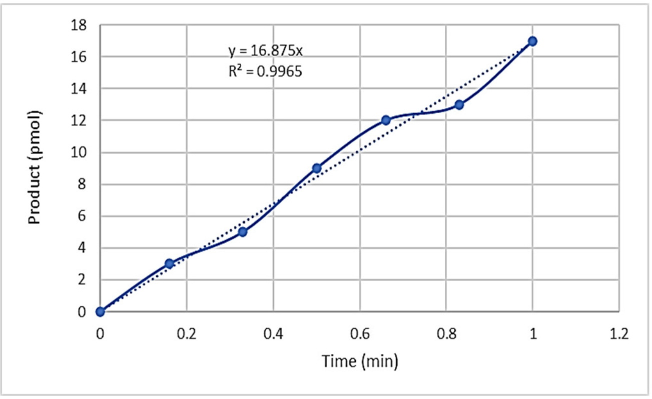


m-TREX1 with 100 nM U3 Unprocessed HIV-1 DNA substrate

(d)

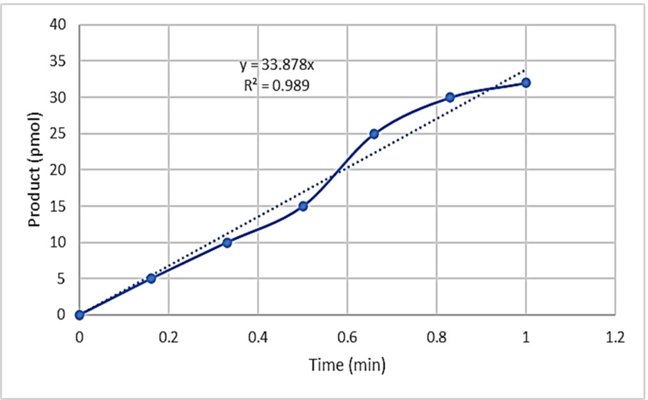


m-TREX1 with 200 nM U3 Unprocessed HIV-1 DNA substrate

(e)

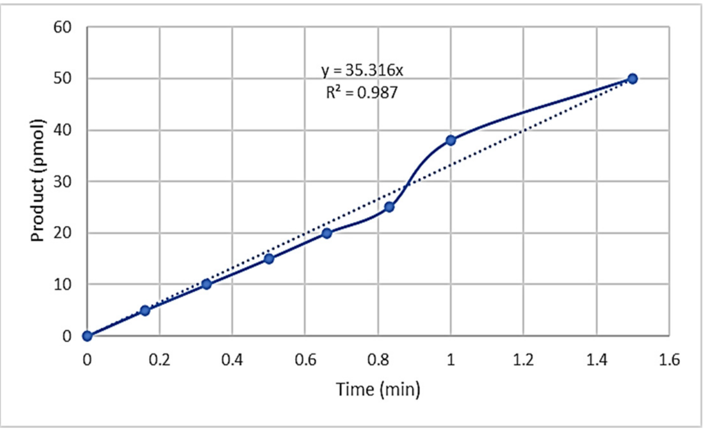


m-TREX1 with 400 nM U3 Unprocessed HIV-1 DNA substrate

(f)

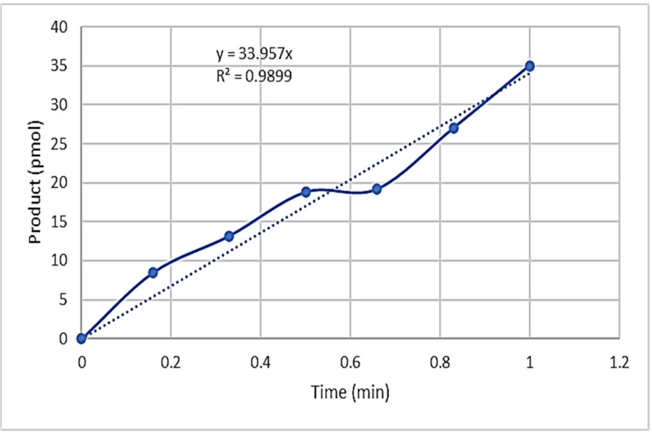


m-TREX1 with 600 nM U3 Unprocessed HIV-1 DNA substrate

### Supplementary Figures

#### Figure 4: Representative Initial velocity curve analysis of m-TREX1 kinetics with U3 processed HIV-1 substrates

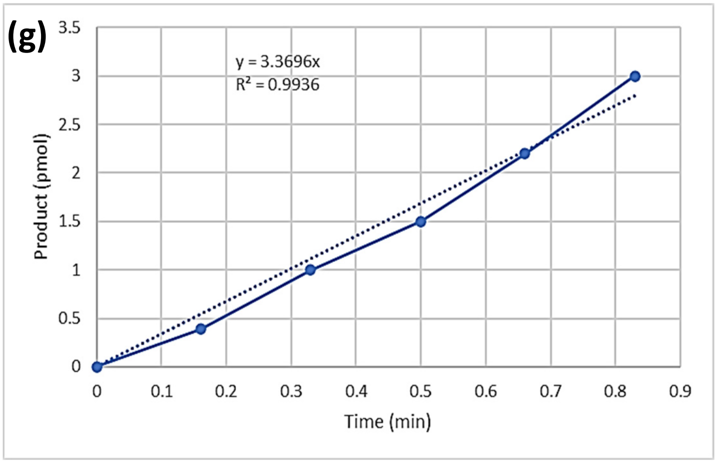


**m-TREX1 with 25 nM U3 Processed HIV-1 DNA substrate**

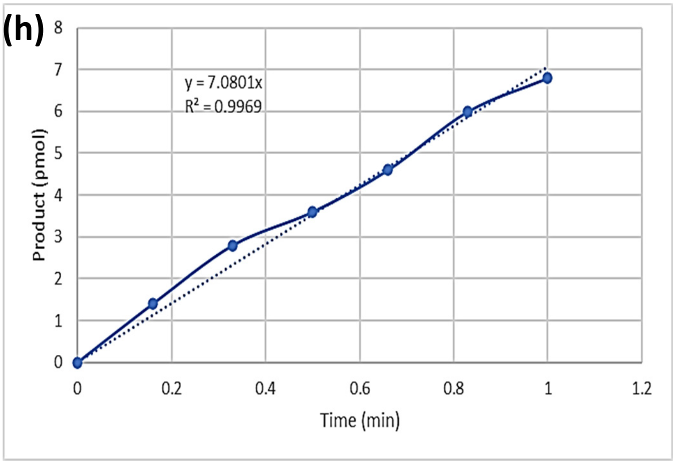


**m-TREX1 with 50 nM U3 Processed HIV-1 DNA substrate**

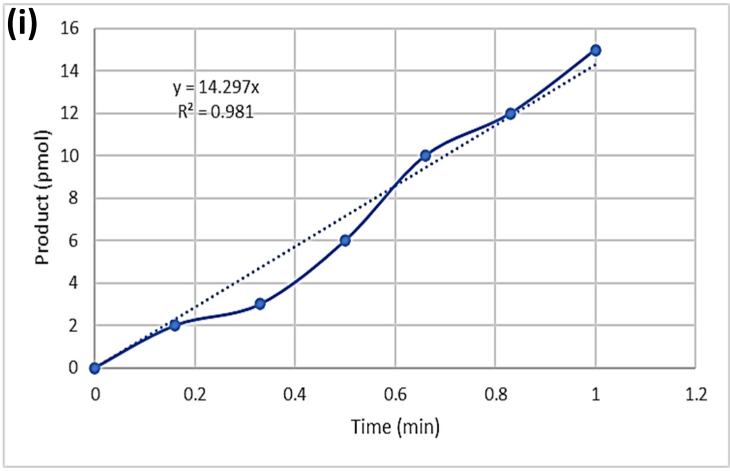


**m-TREX1 with 100 nM U3 Processed HIV-1 DNA substrate**

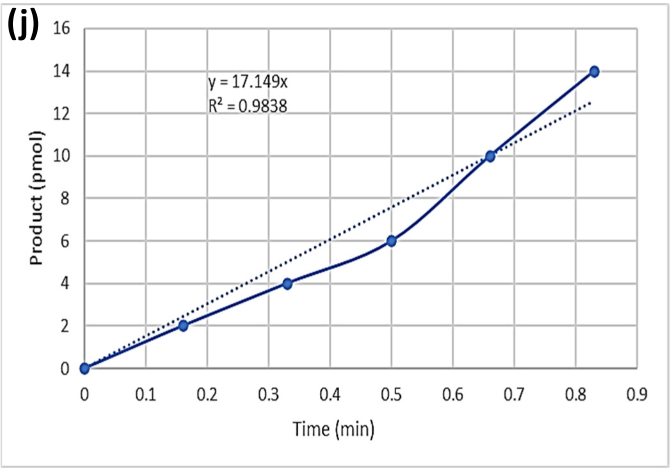


**m-TREX1 with 200 nM U3 Processed HIV-1 DNA substrate**

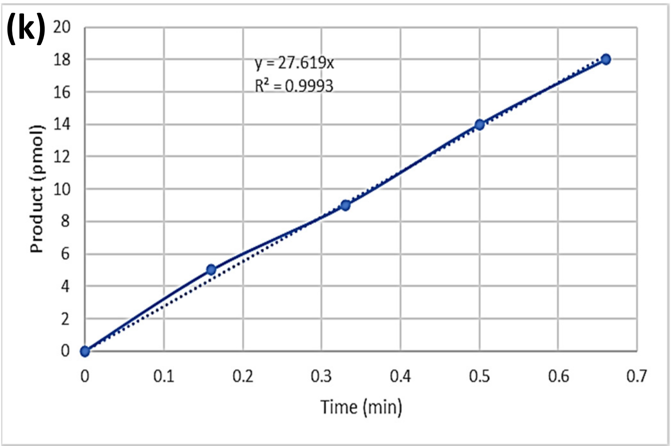


**m-TREX1 with 400 nM U3 Processed HIV-1 DNA substrate**
